# Why is Cyclin A degraded before Cyclin B?

**DOI:** 10.64898/2026.10.23.761299

**Authors:** Annalisa Roberti, Sofia Garcia Soto, Ynon Weiss, Theodoros I. Roumeliotis, Jyoti S. Choudhary, Jonathon Pines

## Abstract

The Anaphase Promoting Complex/Cyclosome (APC/C) ubiquitin ligase controls mitosis by targeting the mitotic Cyclins for degradation. CyclinA has evolved multiple APC/C degrons that enable its degradation as soon as cells enter mitosis while the Spindle Assembly Checkpoint prevents recognition of CyclinB until all chromosomes have attached to the mitotic apparatus. But why the rapid early degradation of CyclinA is conserved in evolution is not known. Here, we mutated two APC/C-recognition motifs in human CyclinA2 to slow down its degradation. We find that slower degradation of CyclinA induces errors in chromosome congression and chromosome clustering at mitotic exit, leading to aneuploidy and malformed daughter cell nuclei. We identify Ki67 as the candidate that interferes with chromosome clustering since its phosphorylation and behaviour are perturbed when CyclinA is degraded more slowly, and increased microtubule dynamics as a contributing factor to chromosome congression errors. We conclude that rapid early CyclinA degradation is important for genomic stability.

## Introduction

The function of mitosis is to segregate an equal and identical set of chromosomes to two daughter cells. To achieve this, the attachment of sister chromatids to the mitotic apparatus is coupled to the mechanism that controls the timing of their separation^1–4^. At the heart of the control mechanism is the Anaphase Promoting Complex/Cyclosome ubiquitin ligase that targets for degradation the two inhibitors of sister chromatid segregation: CyclinB and Securin^5–7^. Sister chromatid attachment is coupled to control of the APC/C by the Spindle Assembly Checkpoint (SAC). The SAC is activated by unattached kinetochores and produces an inhibitor, the Mitotic Checkpoint Complex (MCC), that prevents the APC/C from recognising CyclinB and Securin^8–10^. Thus, although the APC/C is activated upon entry to mitosis it is kept in check by the SAC until all sister chromatids have attached to the mitotic apparatus, thereby ensuring that sister chromatids will be segregated to opposite spindle poles when CyclinB1 and Securin are degraded.

CyclinA and CyclinB coordinate to initiate mitosis and effect the dramatic rearrangement of subcellular architecture required for the cell to segregate its genome. CyclinA also has a role in promoting DNA replication as well as subsequently driving cells into mitosis^11–16^. Despite the careful control of the APC/C by the SAC, CyclinA has evolved multiple APC/C degrons that enable it to compete with the MCC for the APC/C co-activator, CDC20 ^9,17–23^. Thus, CyclinA is degraded before CyclinB in animal cells ^11,24–30^. This raises the question of why CyclinA evolved to be degraded as soon as cells enter mitosis?

The answer could be to ensure genomic stability. Strongly overexpressing wild type CyclinA or expressing a non-degradable form of CyclinA interferes with chromosome alignment and delays sister chromatid separation ^17,18,31,32^; indeed, CyclinA degradation has been suggested to act as a timer in prometaphase that enables proper error correction ^31^ because there is a linear relation between the level of over-expressed CyclinA and the duration of prometaphase^17^. Duane Compton and colleagues showed that increasing the level of CyclinA promotes kinetochore microtubule instability, which delays the stable attachment of kinetochores to microtubules^31–33^. They further showed that depleting CyclinA induces more stable microtubules and promotes stable kinetochore-microtubule attachment. They concluded that CyclinA is degraded in prometaphase to induce a change from the dynamic kinetochore microtubules that aid initial kinetochore capture and spindle assembly to more stable kinetochore microtubules required to maintain kinetochore attachment and ensure proper sister chromatid segregation ^31–33^.

While insightful, these studies overexpressed ectopic CyclinA, and overexpressing Cyclins can perturb substrate specificity due to the plasticity of Cyclin-CDK binding to substrates^34,35^; indeed, in some cell types, Cyclin A or CyclinB can be eliminated, which implies that they can compensate for each other. Thus, the question of why CyclinA evolved to be degraded before CyclinB when the SAC is active is unresolved.

Here, we have mutated the APC/C recognition motifs in endogenous CyclinA to engineer cells in which CyclinA degradation begins at the correct time but proceeds at a slower rate than normal. We find that rapid early CyclinA degradation is important because cells in which CyclinA is degraded more slowly have an increased incidence of chromosome mis-segregation correlating with increased microtubule dynamics, and are less fit compared to control cells. Moreover, slower CyclinA degradation caused problems in mitotic exit, including cells with multiple small nuclei. To elucidate the mechanism responsible, we used mass spectrometry-based phosphoproteomics to identify those proteins whose mitotic phosphorylation was altered when compared to control cells and identified the ‘chromosome surfactant’ Ki67. We found that slower degradation of CyclinA correlated with an increase in the amount of Ki67 on anaphase chromosomes, which could perturb chromosome clustering. We conclude that the rapid loss of CyclinA immediately after nuclear envelope breakdown (NEBD) is important for genomic stability and accurate mitosis.

## Results and Discussion

### The ABBA motif and the second Destruction box synergise to promote CyclinA degradation

To measure the rate of CyclinA degradation we used the ‘nickase’ mutant of CRISPR/Cas9^36^ to introduce an mEmerald tag at the C-terminus of CyclinA, and an emiRFP670 tag ^37^ into the PCNA locus to give a live-cell readout of the cell cycle phase ^38^ in diploid, non-transformed RPE-1 cells. In agreement with previous studies, neither the mEmerald tag on CyclinA^39^ nor the emiRFP tag on PCNA ^38^ perturbed cell cycle or mitotic timing, nor the SAC nor genomic stability (Supplementary Material Fig. S1).

To determine the physiological role of the prometaphase degradation of CyclinA, we mutated APC/C recognition motifs in the amino terminus of CyclinA2. Human CyclinA2 has four APC/C recognition motifs: two ‘Destruction boxes’ (D-box)^40^, one ABBA motif^21^, and one KEN box^41^. Of these, the second D-box (D2box) was the most critical motif for ubiquitylation in vitro and for the degradation of ectopically expressed CyclinA2 in vivo, but synergised with the KEN box, ABBA motif and the Cks1 subunit of the complex^23^. We sought to slow down but not prevent CyclinA degradation; therefore, we made alanine substitutions in the ABBA motif (FTIHVDE) and the essential leucine of the D2 box, singly and together (Fig. 1A). Homozygous CyclinA2^ΔABBA^, CyclinA2^ΔD2box^, and CyclinA2^D2boxΔABB^ mutants were identified by PCR and confirmed by genomic sequencing, RT-PCR and western blot analysis (see Materials and Methods and Supplementary Fig, S2 A-D).

**Figure 1.**
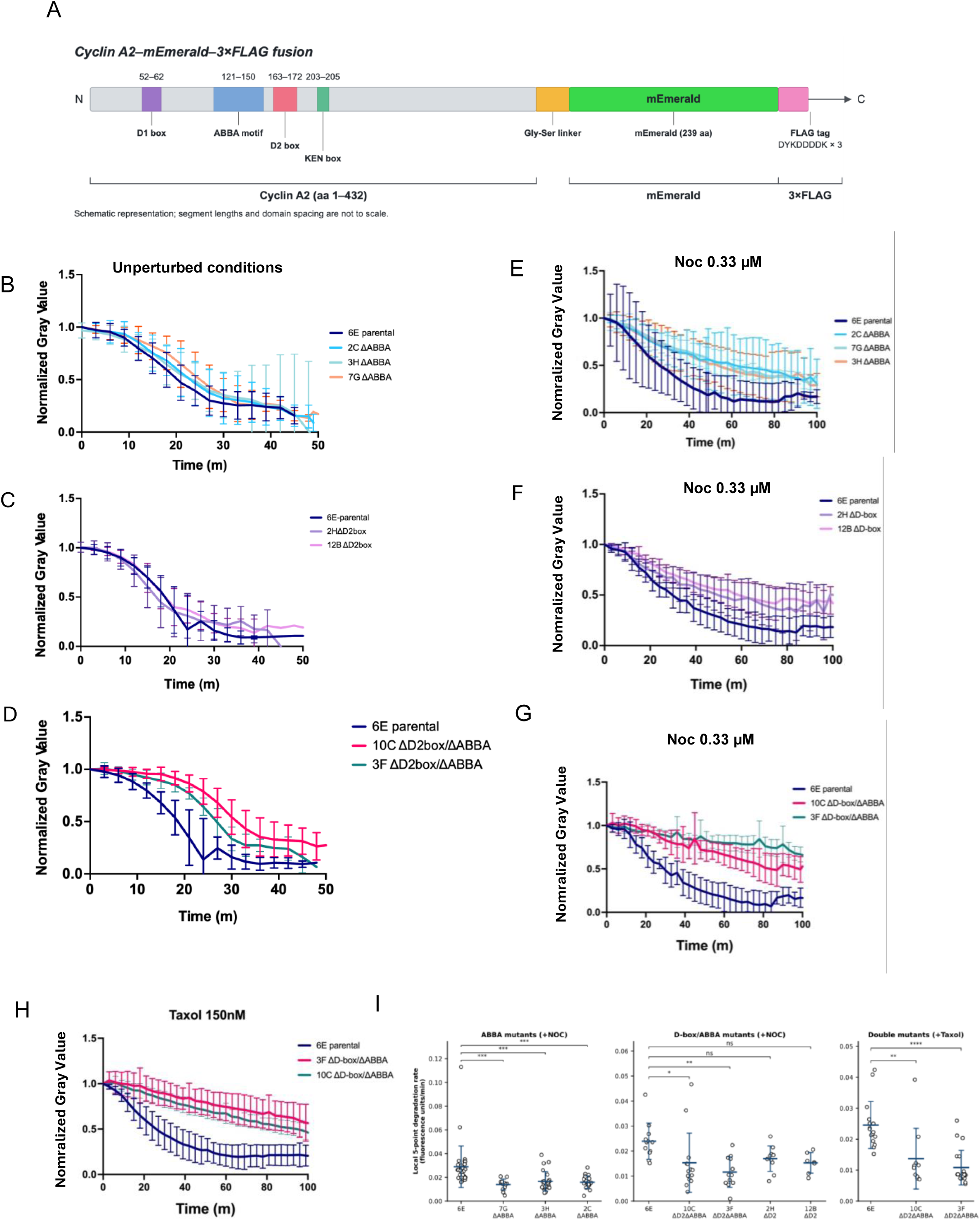
The ABBA and D2box motifs of CyclinA2 cooperate to enable degradation when the SAC is active. (A) Cartoon of Cyclin A-mEmerald fusion protein showing the position of the degrons, linker, the mEmerald and Flag tags. (B-D) CyclinA2 degradation curves in ΔABBA (B), ΔD2box (C) and ΔD2box/ΔABBA(D) clones compared to the parental Cyclin A2-mEmerald cells; time 0 = NEBD. Note that the ΔD2box and the ΔD2box/ΔABBAmutants were filmed at the time, whereas as the ΔABBA was filmed separately. Hence the parental control is the same for the ΔD2box and the ΔD2box/ΔABBAmutants. (E-F) CyclinA2 degradation curves in nocodazole-treated cells of ΔABBA (E), ΔD2-box (F) and ΔD2boxΔABBA (G) clones compared to the parental Cyclin A2-mEmerald cells. Cells were treated with 0.33 µM nocodazole and time 0 = NEBD. (H) CyclinA2 degradation curves in ΔD2boxΔABBA clones in 150 nM paclitaxel and time 0 = NEBD. For all graphs, CyclinA2–mEmerald fluorescence was normalized to its value at NEBD. Plots show mean ± SD. For each condition, a minimum of 10 cells were analysed from at least two independent experiments. Statistical significance was assessed separately within each experiment using one-way ANOVA followed by Dunnett’s multiple-comparisons test, comparing each mutant clone with its matched parental CyclinA2-mEmerald control under the same treatment condition. Full statistical results and adjusted P values are provided in Supplementary Fig. S3. (I) Maximum local Cyclin A2 degradation rates in the indicated clones treated with 0.33 µM nocodazole or 150 nM taxol, calculated using a five-point linear fit. Each dot represents one cell; data are mean ± SD from 2 independent experiments.

We used time-lapse fluorescence and DIC microscopy to assay the timing and rate of CyclinA2 degradation in mitosis in three CyclinA2^ΔABBA^ clones, two CyclinA2^ΔD2box^ clones and two CyclinA2 ^D2boxΔABBA^ clones. As expected, parental wild type CyclinA2-mEmerald began to be degraded at NEBD and was mostly complete after 30 minutes (Fig. 1B). Its degradation rate was reduced when cells were treated with 0.33 µM nocodazole to destabilise microtubules and maintain the SAC at full strength (Fig. 1E). Unexpectedly, mutating the D2box alone did not significantly affect the rate at which CyclinA2 was degraded in an unperturbed mitosis (Fig. 1C), in contrast to its ectopic expression^23^, but this mutant was degraded more slowly in cells treated with nocodazole (Fig. 1F). By comparison, mutating the ABBA motif did slow down the degradation of CyclinA2 in unperturbed mitosis as well as nocodazole-treated cells (Fig. 1B & E). In agreement with in vitro ubiquitylation studies we found that the ABBA and D2box motifs cooperate because the CyclinA2 ^D2boxΔABBA^ double mutant was degraded markedly more slowly than the single mutants (Fig. 1D). In untreated cells, although the maximum degradation rate was not significantly altered in the CyclinA2 ^ΔABBA^ and CyclinA2^ΔABBA/D2box^ mutant clones, it was reached significantly later compared with parental cells, consistent with the mutants being less able to compete with the MCC for APC/C^CDC20^ (Suppl. Fig. S3 A-C). In agreement with this, the local five-point degradation rate was markedly reduced in nocodazole-treated mutant cells (Fig. 1E–G). In the ΔABBA experiment, the mean maximum local degradation rate, calculated using a five-point linear fit for each cell, decreased from 0.025 fluorescence units min⁻¹ in parental Cyclin A2–mEmerald cells to 0.014–0.016 fluorescence units min⁻¹ in the Cyclin A2 ΔABBA clones. In the double-mutant experiment, the mean maximum local degradation rate decreased from 0.024 fluorescence units min⁻¹ in parental cells to 0.013 and 0.010 fluorescence units min⁻¹ in clones 10C and 3F, respectively. A similar reduction was observed in taxol-treated double-mutant cells, with mean rates of 0.014 and 0.010 fluorescence units min⁻¹ in clones 10C and 3F, respectively, compared with 0.025 fluorescence units min⁻¹ in parental cells (Fig. 1H & I). Consistent with their reduced degradation rates, the mutant cells retained more Cyclin A2 at 21 min after NEBD. Under nocodazole treatment, parental cells had degraded approximately 43% of Cyclin A2, compared with 20–24% in the ΔABBA clones. In the double-mutant experiment, parental cells had degraded approximately 36% of Cyclin A2, whereas only 6–9% had been degraded in the ΔD2-boxΔABBA clones. Although the local five-point degradation rate was not consistently reduced in the ΔD2-box clones, these cells had degraded only 20–26% of Cyclin A2 at 21 min, compared with approximately 36% in the corresponding parental cells. These results support a model in which the ABBA and D2-box motifs cooperate to enable Cyclin A2 to compete with the MCC for APC/C^CDC20^ binding and thereby promote its rapid degradation during prometaphase. (Suppl. Fig. S3 D & E).

### Slower CyclinA degradation destabilises kinetochore-microtubule attachment

Overexpressing CyclinA2 or ectopic expression of a non-degradable CyclinA2 mutant interferes with chromosome congression and prolongs prometaphase^17,18,31^, at least in part because microtubules are made more dynamic ^31–33^, but the question remained whether CyclinA at endogenous levels could also delay prometaphase. Time-lapse analysis showed that slowing down the degradation of endogenous CyclinA did indeed increase the time between NEBD and anaphase for both the CyclinA2^ΔABBA^ and the CyclinA2 ^D2boxΔABBA^ mutant cell lines (Fig. 2A & B). This prompted us to measure the stability of microtubules in parental and CyclinA2 mutant prometaphase cells. We used SPY555-DNA to visualise chromosomes and SiR-tubulin to mark the microtubules. Both the CyclinA2^ΔABBA^ single mutant and the CyclinA2 ^D2boxΔABBA^ mutant showed faster fluorescence loss and reduced area under the curve compared to parental CyclinA2-mEmerald cells, consistent with increased microtubule instability and an extended prometaphase (Fig. 2C & D). As a second read-out for microtubule stability, we measured the length of prometaphase spindles -as marked by MAD1 staining at kinetochores - and metaphase spindles. Consistent with reduced MT stability, prometaphase spindles were shorter in the two mutant cell lines (Fig. 2E & F).

**Figure 2.**
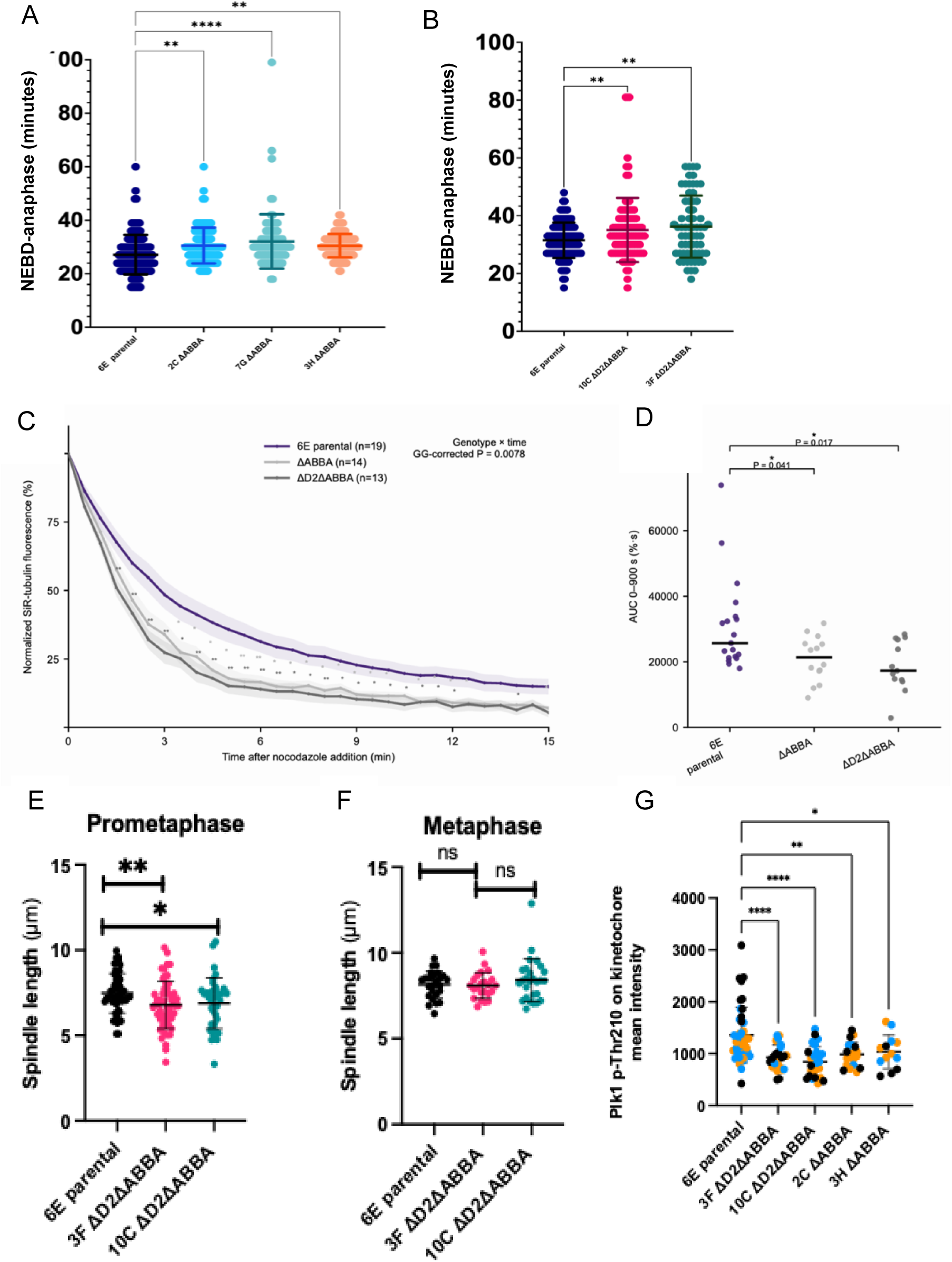
Slower CyclinA degradation prolongs the period of microtubule instability. (A-B) Mitotic timing (NEBD to anaphase) measured by time-lapse DIC microscopy in parental CyclinA2-mEmerald and mutant cell lines (CyclinA2^ΔABBA^ and CyclinA2^ΔD2box/ΔABBA^). (C) SiR-tubulin fluorescence intensity following treatment with 0.66 µM nocodazole normalized to the initial value (t = 0, 100%). Curves show mean ± SEM. Statistical significance was assessed over 30s–15 min using a two-way repeated-measures ANOVA with Greenhouse–Geisser correction. **P < 0.01 for the genotype × time interaction (P = 0.0086). (D) Area under the observed fluorescence-decay curve (AUC). Values were compared using Welch’s ANOVA (P = 0.014). Data were pooled from three independent experiments: parental CyclinA2-mEmerald 6E n = 19 cells; ΔABBA n = 14 cells; ΔD2box/ΔABBA n = 13 cells. Curves were compared over 30s–15 min using a two-way repeated-measures ANOVA with Greenhouse–Geisser correction (genotype × time interaction, P = 0.0086). AUC values were compared using Welch’s ANOVA (P = 0.014). (E-F) Quantification of spindle length in prometaphase (E) and metaphase (F) cells in parental CyclinA2-mEmerald and CyclinA2ΔD2box/ΔABBA. (G) PLK1 phospho-T210 staining at kinetochores. Background-corrected mean fluorescence intensity in parental CyclinA2-mEmerald 6E and CyclinA2 mutant cells. One-way ANOVA with Dunnett’s multiple-comparisons test versus 6E. *P < 0.05, **P < 0.01, ****P < 0.0001.

We conclude that the rate of CyclinA2 degradation acts as a timer in mitosis because reducing it results in a prolonged period of microtubule instability that extends prometaphase.

### Slower CyclinA degradation causes errors in both chromosome segregation and the formation of daughter cells

Our time-lapse analysis revealed that slowing down CyclinA2 degradation increased the number of mitotic errors. The parental CyclinA2-mEmerald RPE1 cells had an error rate of ≤1% across all chromosome error categories, whereas for the CyclinA2^ΔABBA^ mutant clones 15-18% had chromatin bridges, 4-6% had problems in chromosome congression, 3-6% had persistent polar chromosomes and 2-4% had lagging chromosomes (Fig. 3A & C). The CyclinA2^ΔD2box/ΔABBA^ clones had similar error profiles but slightly higher frequencies of errors (Fig. 3A & C). The frequency of micronuclei was higher in both CyclinA^ΔD2box/ΔABBA^ (7.96%) and CyclinA2^ΔABBA^ mutant clones (10.39%) compared to CyclinA2-mEmerald parental cells (5.14%) (Fig. 3D and Supplementary Fig.3A). Consistent with the increased chromosome segregation defects, metaphase spreads revealed increased aneuploidy in the CyclinA2 mutant cells. The CyclinA2^ΔD2box/ΔABBA^ mutants also exhibited a small population of polyploid cells, which likely accounted for their increased nuclear area compared with the parental CyclinA2-mEmerald and CyclinA2^ΔABBA^ cells (Supplementary Fig. S4 A-C).

**Figure 3.**
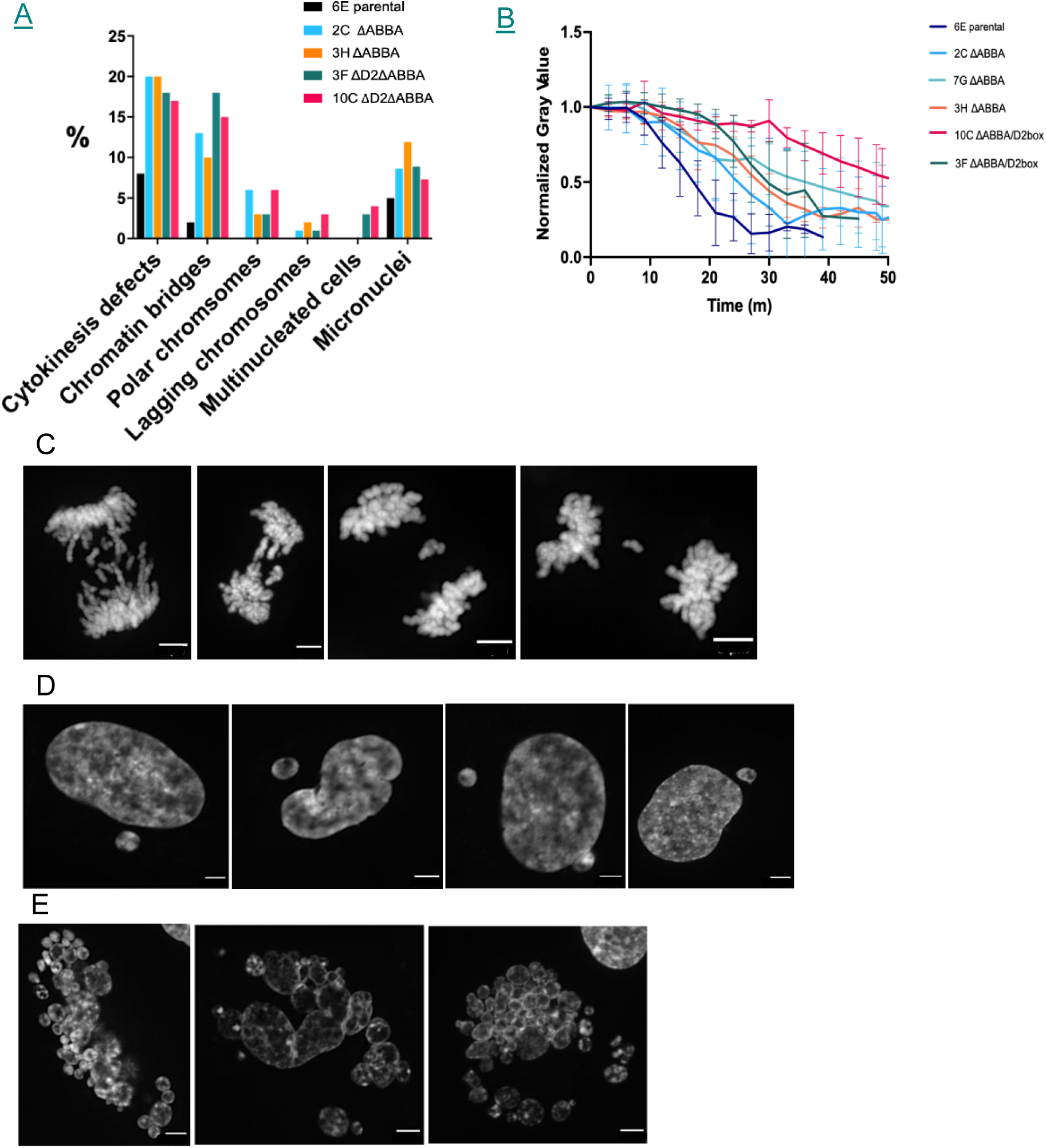
Slower CyclinA degradation correlates with increased errors in mitosis. (A) Percentage of mitotic defects in parental CyclinA2-mEmerald and mutant cell lines (CyclinA2^ΔABBA^ and CyclinA2^ΔD2box/ΔABBA^). (B) CyclinA2-mEmerald degradation over time in those CyclinA2^ΔABBA^ and CyclinA2^ΔD2box/ΔABBA^ cells exhibiting mitotic defects, compared to parental cells. (C) Representative examples of lagging chromosomes in ΔABBA and ΔD2boxΔABBA cells. (D) Representative examples of micronuclei in ΔABBA and ΔD2boxΔABBA cells. (E) Representative examples of multi-nucleated cells observed in ΔD2boxΔABBA cells. Cells stained with DAPI. Scale bar, 5 μm.

When we measured the degradation of CyclinA2-mEmerald specifically in those cells that had subsequently exhibited mitotic errors and compared this to parental cells we found that there was a strong correlation between slower CyclinA-mEmerald degradation and errors in mitosis (Fig. 3B).

These errors were consistent with perturbed kinetochore-microtubule attachments induced by increased microtubule instability, but cytokinesis defects and chromatin bridges can also be caused by damaged DNA stemming from problems in DNA replication where CyclinA2 has a role. To test for this, we assayed cells for the levels of DNA damage as visualised by antibodies against γ-H2AX and 53BP1, and DNA replication using CldU and IdU and DNA combing to measure origin firing and the elongation rate of DNA polymerases. Across three independent experiments, there was a small increase in the fraction of γH2AX-positive nuclei in CyclinA2 ^ΔD2box/ΔABBA^ mutant cells, from 57.7 ± 9.3% in parental CyclinA2-mEmerald 6E cells, to 66.7 ± 3.1% and 68.3 ± 8.4% in the CyclinA2^ΔD2box/ΔABBA^ clones 3F and 10C, respectively (Supplementary Fig.S4D), which could be accounted for by the increase in stalled forks from 4% in parental 6E cells to 14% in the 3F mutant (Supplementary Fig. S4E). In contrast, the mean number of γH2AX foci per nucleus was not significantly different between parental and mutant cells, and we did not detect any difference in origin firing or the elongation rate of DNA polymerases.

We conclude that slowing down CyclinA degradation induces an increased rate of mitotic errors largely through effects on mitosis rather than S phase.

### Mass spectrometry identifies prometaphase CyclinA-CDK substrates

To determine the mechanisms by which slower degradation of CyclinA perturbed mitosis, we used mass spectrometry-based phosphoproteomics to identify those proteins whose phosphorylation in prometaphase was altered in the CyclinA2^ΔD2box/ΔABBA^ mutant clones. Cells were enriched in mitosis using 150 nM taxol and collected at a time when CyclinA2 degradation was complete in parental 6E cells but not in the CyclinA2^ΔD2box/ΔABBA^ mutant clones 3F and 10C (Supplementary Fig. S5). We focused on phosphorylation sites that changed concordantly in both mutant clones and whose alteration was not explained by a corresponding change in the total protein abundance. This identified 30 sites on 30 proteins that were consistently hyperphosphorylated and 19 sites on 15 proteins that were consistently hypophosphorylated (Supplementary Table S1).

CyclinA–CDK1 has been shown to promote the correction of erroneous kinetochore–microtubule attachments and timely progression through prometaphase^13,27–29^. Previous studies identified potential mitotic CyclinA–CDK substrates using various strategies, including overexpression, degron-mediated depletion, and analysis of cells maintained for several hours in prometaphase to drive complete CyclinA degradation^14,29,39^. When we compared the phosphoproteins identified in our study with those reported previously, we found 19 proteins in common with at least one other study (Table 1).

We interrogated our mass spectrometry data to identify candidate proteins whose persistent phosphorylation could be responsible for the phenotypes we observed in cells when CyclinA degradation was slowed down: increased microtubule dynamics, defective chromosome congression, polar chromosomes, and daughter cells with multiple small nuclei. Among the proteins associated with microtubule dynamics, we found that S592 on MAP1S and S1756 on KIF20B were consistently hyperphosphorylated in both mutant clones despite unchanged or slightly reduced total protein abundance. KIF20B was a particularly compelling candidate because phosphorylation of S1756 was increased by 0.72 and 0.79 log2-fold in CyclinA2^ΔD2box/ΔABBA^3F and 10C clones, respectively, while total KIF20B abundance remained unchanged. Moreover, S1756 comprises part of a SSSK sequence that matches the non-proline-directed CDK1 consensus motif identified by Al-Rawi et al.^42^ as being preferentially phosphorylated by CyclinA–CDK1 and promoted by Cks. However, immunofluorescence analysis did not reveal substantial changes in KIF20B abundance or localisation in the CyclinA2 mutant cells. We were unable to assess whether MAP1S abundance or localisation was changed because the antibodies we tested did not give specific and reliable staining.

Aside from problems in forming stable kinetochore-microtubule attachments due to more dynamic microtubules^31^, the Compton lab reported that defective chromosome congression in CyclinA-overexpressing cells can also be caused by perturbing MYPT1 localisation to kinetochores to counteract PLK1 kinase activity^33^. Although we detected MYPT1 in our phosphoproteome dataset, its phosphorylation was not altered consistently in both CyclinA2^ΔD2box/ΔABBA^ double mutant clones; therefore, we assessed whether kinetochore-associated PLK1 activity was changed by measuring the phosphorylation of PLK1 at its activating T210 residue. Our mass spectrometry data did not show any global change at this site, but when we stained cells with a phospho-specific antibody, this revealed that the signal for phospho-T210 was reduced at the kinetochores of the CyclinA2^ΔD2box/ΔABBA^ mutant cells compared to parental cells (Fig. 2F). This indicated that Plk1 was less active when CyclinA degradation was slower, just as in cells over-expressing CyclinA^33^. This could result in more unstable kinetochore-microtubule attachment and lead to increased errors in chromosome segregation.

### Ki67 abundance on chromosomes is altered when CyclinA degradation is slow

We were intrigued by our observation that some of the daughter cells of the CyclinA2^ΔD2box/ΔABBA^ mutants formed multiple small nuclei (3% of 3F cells and 4% of 10C cells) (Fig. 3A and Fig. 3E), a phenotype that was not observed in either parental cells or the single ΔABBA mutants.

This phenotype resembled perturbation of the ‘surfactant’ property of the chromosome periphery that causes chromosomes to remain as individuals rather than clustering together when reforming the nuclear envelope^43,44^. Ki67 is the key ‘molecular surfactant’^43^ that maintains the individuality of chromosomes through electrostatic repulsion and is hyperphosphorylated in mitosis by CDK1 ^43,44^ (reviewed in ref. ^45^). We noted that our mass spectrometry analysis displayed site-specific changes in phosphorylation of Ki67 in the CyclinA2 ^ΔD2boxΔABBA^ mutant clones: residues S1861/S1864 were hyperphosphorylated despite essentially unchanged total Ki67 abundance. S1861 had been previously identified as a potential CyclinA–CDK1-regulated site^29^ and Ki67 was identified as a protein containing CyclinA- and Cks1-dependent phosphosites^38^, although the sites reported in that study were different from those identified here. We reasoned that Ki67 localisation might be perturbed in cells when CyclinA degradation is slowed down; therefore, we analysed the behaviour of Ki67 by immunofluorescence. This showed a marked difference between the CyclinA2-mEmerald parental and CyclinA2^ΔD2box/ΔABBA^ mutant cells in Ki67 abundance on chromosomes in mitosis. Ki67 protein accumulates through the cell cycle^46^ and we found there was more Ki67 signal at the chromosome periphery in the CyclinA2^ΔD2box/ΔABBA^mutant clones, and the ratio of chromosomal to cytoplasmic Ki67 signal was markedly increased. This difference was also observed in prometaphase cells arrested with DMA (Fig. 4A-C). The difference in Ki67 recruitment to chromosomes between wild type and CyclinA2^ΔD2boxΔABBA^ cells was even more marked in anaphase (Fig. 4D), where Ki67 would be predicted to inhibit the chromosome clustering required to generate a single nucleus encompassing all 46 chromosomes in each daughter cell. Inhibiting chromosome clustering has been shown to lead to multiple small nuclei forming around individual or small groups of chromosomes^43,44^, just as we saw in the CyclinA2 ^ΔD2boxΔABBA^cells. The CyclinA2^ΔABBA^ mutant did not show any difference in Ki67 recruitment from the parental cells, consistent with the absence of detectable multi-nucleated cells in this population.

**Figure 4.**
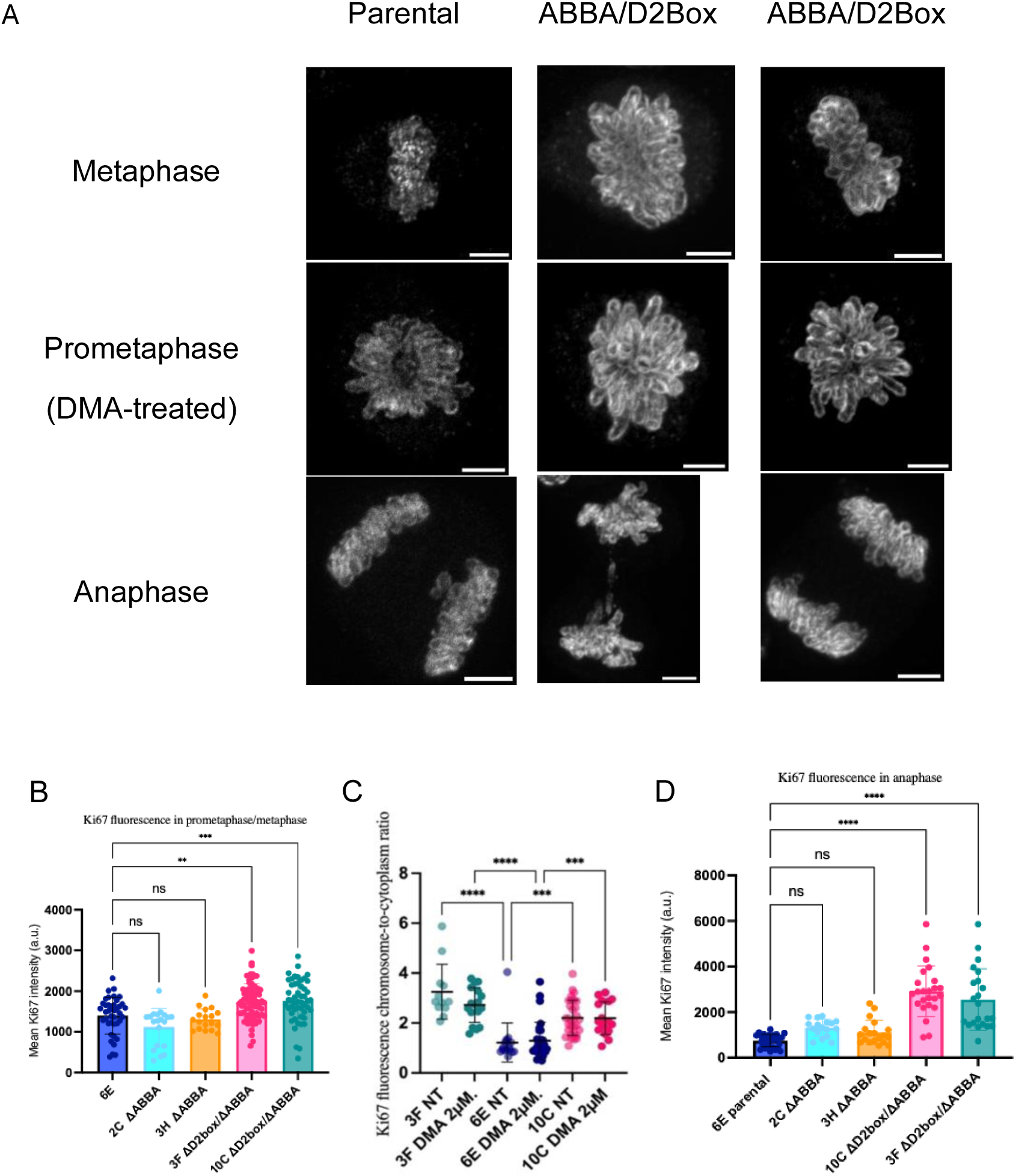
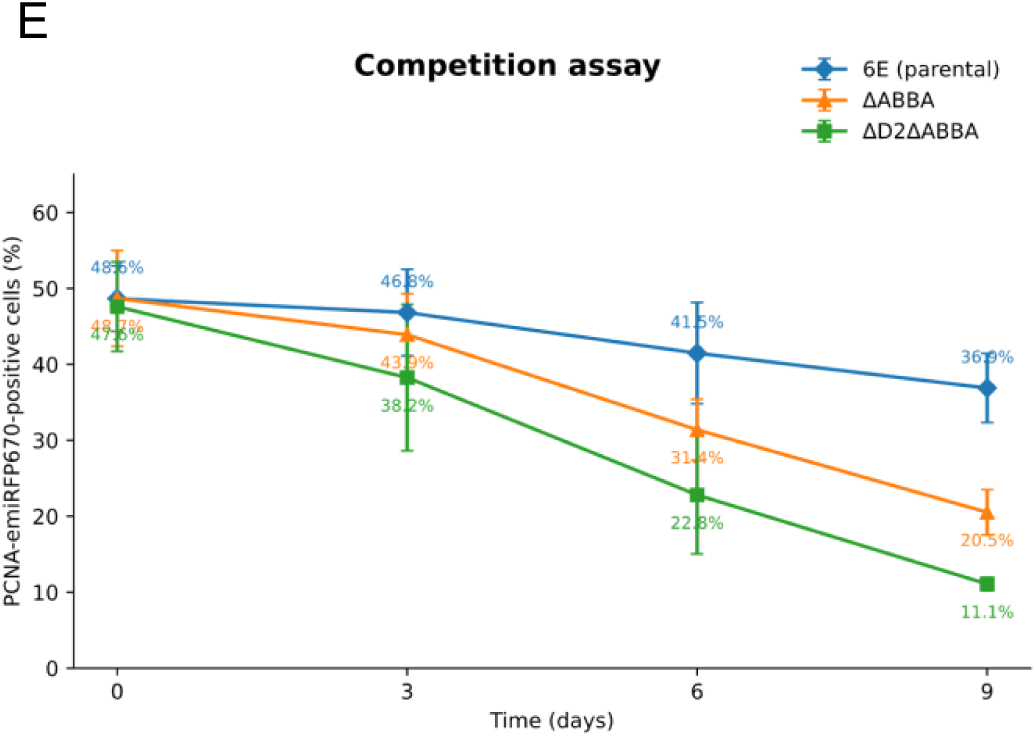
CyclinA2^ΔABBA/D2box^ mutants are less fit and display altered Ki67 abundance and chromosome association in mitosis. (A) Representative images of Ki67 fluorescence in parental CyclinA2-mEmerald (left panels) and CyclinA2 ^D2boxΔABBA^ mutant cells (middle and right panels) during prometaphase, metaphase and anaphase. The scale bar corresponds to 5 μm. (B) Quantification of Ki67 fluorescence intensity in prometaphase/metaphase cells. Individual points represent single cells; horizontal lines indicate the mean ± SD (C) Ki67 chromosome-to-cytoplasm fluorescence ratio in untreated (NT) and DMA-treated cells. Individual points represent single cells; mean ± SD is shown. (D) Quantification of chromosomal Ki67 fluorescence intensity in anaphase cells. Individual points represent single cells; mean ± SD is shown. (E) Competition assay showing the proportion of CyclinA2-mEmerald-positive cells over time in parental and Cyclin A mutant cell populations. Data are presented as mean ± SD from two independent experiments, combining two independent clones for each mutant genotype (ΔABBA: 2C and 3H; ΔD2box/ΔABBA: 10C and 3F). Two-way RM ANOVA, genotype × time: P < 0.0001.

### Cells that degrade CyclinA more slowly are less fit

The conservation through evolution of the early mitotic degradation of CyclinA indicated that cells where the rate of CyclinA degradation is reduced could be at a disadvantage compared to wild type cells. To test this, we set up a competition assay between unmodified RPE1-hTert cells and our wild type and mutant CyclinA-mEmerald RPE1 cell lines. We used the endogenously tagged PCNA as a cell marker for the wild type and mutant CyclinA-mEmerald cell lines. We co-cultured equal quantities of unmodified RPE1 and the CyclinA-tagged cells for one week and assayed the proportion of the two cells in the culture by flow cytometry to detect PCNA. We found that over 9 days there was a small change in the 50:50 ratio of unmodified RPE1 to cells with wild type CyclinA-mEmerald, but the ratio of unmodified RPE1 to either of the CyclinA mutant cell lines rapidly changed such that the CyclinA mutant cells became the minority population (Fig. 4E).

To determine whether the mitotic defects observed in CyclinA mutant cells could account for their progressive depletion from the proliferating population, we modelled the competition assay on a per-cell-division basis. We assumed that following each division a fraction of the mutant progeny would fail to contribute to subsequent rounds of proliferation. Population propagation was therefore described as M_n+1_=2M_n_(1-x), where M_n_ represents the proliferating mutant population after n divisions and x represents the effective fractional loss of proliferative capacity per cell division. The CyclinA2-mEmerald parental (6E) cells were used as the reference, and the number of cell divisions was estimated using a doubling time of approximately 18 h.

Both CyclinA2^ΔABBA^ and CyclinA2^ΔD2box/ΔABBA^ cells were progressively depleted relative to unmodified RPE1 cells during the competition assay, and the effect was considerably stronger for the CyclinA2^ΔD2box/ΔABBA^ mutants. Modelling the complete competition time course indicated that the behaviour of the ΔABBA clones could be reproduced by an additional effective loss of approximately 3–4% of proliferative capacity per cell division relative to parental cells. In contrast, the progressive depletion of the ΔD2boxΔABBA clones required an additional loss of approximately 10 –11% per division.

We then asked whether the magnitude of this loss was compatible with the observed frequency of mitotic defects. Cytokinesis defects increased from approximately 8% in parental cells to approximately 20% in the CyclinA2^ΔABBA^ clones and 17–18% in the CyclinA2^ΔD2box/ΔABBA^ clones. Chromatin bridges increased from approximately 2% in parental cells to 10–15% in CyclinA2^ΔABBA^ cells and 15–18% in CyclinA2^ΔD2box/ΔABBA^ cells, while lagging chromosomes occurred at lower frequencies. Because individual cells could display more than one mitotic phenotype, these categories were not considered additive. Multinucleated cells and micronuclei were considered downstream outcomes of defective mitoses rather than independent mitotic defects and were not included as additional events in the population model. The observed frequency of mitotic defects was sufficient to account for the loss predicted by the model. For CyclinA2^ΔABBA^ cells, the predicted loss was substantially lower than the observed frequency of mitotic defects, whereas in CyclinA2^ΔD2box/ΔABBA^ cells the predicted loss was closer to the frequency of the observed defects. Since the frequency of mitotic defects alone did not explain the difference between the two classes of mutants, this indicated that the long-term proliferative consequences of defective mitoses, rather than their frequency alone, may contribute to the different behaviour of the mutants. The consequences of defective mitoses were evident in the resulting progeny. Both CyclinA2^ΔABBA^ and CyclinA2^ΔD2box/ΔABBA^ mutants displayed increased aneuploidy, while the CyclinA2^ΔD2box/ΔABBA^ clones showed an increase in polyploidy, increased nuclear area and multinucleation. Together, these phenotypes indicate that defective mitoses, particularly in the CyclinA2^ΔD2box/ΔABBA^ mutants, can generate progeny with altered chromosomal and nuclear characteristics that compromise their subsequent proliferative capacity. The increased fraction of γH2AX-positive cells observed in the mutant CyclinA clones over the parental cells may represent an additional consequence associated with their reduced proliferative capacity.

Our finding that cells that degrade CyclinA more slowly are less fit and are quickly out-competed by cells that degrade CyclinA with normal kinetics provides a likely explanation for the conservation of the rapid onset of CyclinA degradation in mitosis through animal cell evolution^11,24–30^. Our results show that the rate of CyclinA degradation does act as a timer for prometaphase^17,31^: in agreement with previous reports^31–33^, we identified increased kinetochore microtubule dynamics in prometaphase and a reduction in phospho-Plk1 at kinetochores as important parameters induced by persisting CyclinA. The effect on mitotic fidelity, however, depends on cellular context: overexpressing CyclinA reduced the frequency of lagging chromosomes in chromosomally unstable U2OS cancer cells^31^, whereas we found that slower CyclinA degradation increased mitotic errors in diploid, non-transformed RPE1 cells. Moreover, we have uncovered the necessity of early CyclinA degradation for processes beyond microtubule stability; notably for the proper function of the ‘chromosomal surfactant’ Ki67. When CyclinA is degraded too slowly, higher levels of Ki67 associate with anaphase chromosomes, which likely explains the subsequent appearance of daughter cells with multiple small nuclei. These cells should subsequently be eliminated from the population because they should trigger a p53-dependent arrest^47^. In this respect, it will be interesting to study human ES cells where Cyclin A2 is reported to be only partially degraded^48^ (although the evidence for this is based on immunoblots of synchronised cells that lack the accuracy of our single cell assays).

We conclude that the conservation through evolution of the degradation of CyclinA before CyclinB is because rapid CyclinA degradation in early mitosis is required to prevent CyclinA interfering with chromosome congression and perturbing the subsequent formation of daughter cell nuclei.

## Materials and Methods

### Cell culture and drug treatment

hTERT RPE-1 FRT/TO cells were cultured in F12/DMEM medium (Sigma-Aldrich) supplemented with GlutaMAX (Invitrogen), 10% FBS (Gibco), 0.35% sodium bicarbonate, penicillin (100 U/ml), streptomycin (100 µg/ml) and Fungizone (0.5 µ/ml). Cells were maintained in a humidified incubator at 37 °C and 5% CO2 concentration. For live-cell imaging experiments cells were imaged in Leibovitz L-15 medium (Thermo Fisher) supplemented with 10% FBS, penicillin (100 U/ml) and streptomycin (100 µg/ml).

For live cell imaging experiments, cells were seeded in 8-well chamber slides (µSlide, ibidi) and drugs were added directly into the imaging medium prior to acquisition. In the indicated experiments, Cells were stained with 50 nM SiR-DNA or SPY595-DNA (1:20,000 dilution from stock), either alone or in combination with 50 nM SiR-tubulin, following the manufacturer’s protocol or treated with the following drug concentrations: 150 nM paclitaxel (Sigma-Aldrich), DMA (10 µM) and nocodazole (0.33 µM).

### Gene editing

For CCNA2 tagging, 2 million RPE-1 FRT/TO cells were transfected using 1ug of a modified version of the PX466 ‘All-in-One’ plasmid containing Cas9D10A-T2A-mRuby and gRNAs targeting CCNA2 (5’-ATTTACATCTTAGAAAACAA-3’ and 5’-TGGTGTACAGTTTTTAACTT-3’). The all-in-one plasmid was cotransfected with 1ug of repair plasmid designed as a fusion of LINKER-mEmerald flanked by two 500 bp arms, homologous to the genomic region around theCas9 cutting site. All CRISPR repair templates were cloned into the pDrive Amp-plasmid. Seventy-two hours post transfection, 50,000 mEmerald positive cells were sorted in a 1 cm well and expanded for 1 week before a second sorting of single cells in 96-well plates. PCR products of positive clones were sequenced using the same primers. Homozygous integration of the mEmerald tag at the endogenous CCNA2 locus was confirmed by PCR using primers forward 5-ggggcatgcatgtgtgtttc-3 and the reverse 5-TTCACAGAACACAGACCACC-3. PCR products of positive clones were sequenced using the same primers.

RPE1 cells expressing CyclinA2–mEmerald following endogenous knock-in were subsequently infected with rAAV encoding PCNA-emirFP670nano3. Following transduction, far-red-positive cells were single-cell sorted using a BD FACSAria S6 cell sorter into 96-well plates (one cell per well). Individual clones were expanded and screened by high-content microscopy for emirFP670nano3 far-red fluorescence. Positive clones were subsequently characterised by PCR amplification of PCNA was performed using two primer pairs: the first pair comprised forward primer 5′-GCGCAAGCAGGCCTAACTAC-3′ and reverse primer 5′-CGTTTCGATGAGCAAGGC-3′; the second pair comprised forward primer 5′-ACCTTGTATGGAGTCACAGC-3′ and reverse primer 5′-GGATAGCGGTTGCCGAAAT-3′. The same primers were used to sequence the PCR products. To generate CyclinA2 ΔABBA, ΔD2 and ΔABBA/ΔD2 cell lines, RPE-1 CyclinA2-mEmerald^+/+^ FRT/TO cells were transfected using Cas9^D10A^ RiboNucleoProteins (RNPs—Integrated DNA technology). Briefly, sgRNA was assembled by mixing equal volumes of individual gRNA 100 µM with tracr-RNA 100 µM. For production of CyclinA2 ΔABBA, ΔD2 and ΔABBA/ΔD2 cell lines the following gRNAs were used: 5′-GGTCCCTCTTGATTATCCAA -3′ and 5′-TTAGTTTACCTGGACCCAGA-3’. The mixture was incubated at 95 °C for 5 min and then kept at RT for 40 min to anneal sgRNAs. About 4 uL of annealed sgRNA were mixed with 2.5 µL of 62 nM Cas9^D10A^ protein to obtain RNPs. Two million cells were transfected with 18.8 pmol of each RNP together with 6 µg of the corresponding repair plasmid. The repair plasmids contained a “wobbled” second exon of CCNA2 carrying one of the following modifications: an alanine-substituted ABBA motif (PAFTIHVDE), designated ΔABBA; substitution of the D2-box sequence VAPLKDL with AAPAKDA, designated ΔD2; or both modifications, designated ΔD2ΔABBA. Each repair plasmid also contained a neomycin-resistance cassette and was flanked by two 500-bp homology arms corresponding to the genomic regions surrounding the Cas9 cleavage site. Seventy-two hours post transfection, CyclinA2 ΔABBA, ΔD2 and ΔABBA/ΔD2 cells were selected for for 10 days with 0.4 mg/ml Geneticin (Gibco). Single cells were then sorted in 96-well plates. The presence of the ABBA, D2 or double mutations were identified through PCR and sequencing using forward 5′-TATTCTTTGGCCGGGTCGG-3′ and reverse 5′-tggcatggtggttctgttagg3′ primers.

### RT-PCR

Total RNA was extracted from exponentially growing cells using the Monarch Total RNA Miniprep Kit (New England Biolabs), according to the manufacturer’s instructions. 1 µg of total RNA was reverse transcribed into cDNA using SuperScript III Reverse Transcriptase (Invitrogen), following the manufacturer’s instructions. CCNA2 cDNA was amplified using the following primers; all sequences are written 5′→3′.

Forward primers: 5′UTR-F, CTGCTCGCTGCATCTCTGG;

Exon 1-F, TGCTAGCATTGCAGCAGACG;

Exon 2-F, ATGATGAGCATGTCACCGTTCC;

Exon 3-F, AGTCACCACATACTATGGACATG;

Exon 4-F, GTGGGTTACATGAAGAAACAGCC;

Exon 5-F, ACCCCCC CAGAAGTAGCAGAG;

Exon 6-F, GATGCTGACCCATACCTCAAG.

Reverse primers: Exon 4-R, GGCTGTTTCTTCATGTAACCCAC;

Exon 8-R, AGTGTCTCTGGTGGGTTGAG;

3′UTR-R, TTGGCCACAACTTCTGTATTCAG.

For RT–PCR analysis of the CyclinA2 ΔABBA and ΔD2ΔABBA mutant clones, cDNA was amplified using a forward primer specific to the wobble-modified exon 2 sequence (5′-CTGAAAGACCTCCCCGTCAA-3′) and a reverse primer targeting the 3′ UTR, yielding a 2,023-bp amplicon.

### Immunoblotting

Cell lysates were separated through SDS-PAGE on a 4–12% NuPAGE gel (Invitrogen) and transferred to an Immobilon-FL polyvinylidene fluoride membrane (IPFL00010, Millipore). The membrane was blocked with 5% milk, 0.1% Tween in PBS and incubated overnight with primary antibodies at 4 °C in 2.5% milk and 0.1% Tween in PBS. The following day, the membrane was washed with 0.1% Tween in PBS and incubated with secondary antibodies in 2.5% milk and 0.1% Tween in PBS for 1 hour at RT. Membranes were visualised with a LI-COR Odyssey CLx scanner (LI-COR Biosciences). Membranes were probed with a mouse monoclonal anti-CyclinA2 antibody raised against an N-terminal epitope (Moravian Biotechnology; 1:2,000 for WB), a mouse anti-FLAG M2 antibody (1:6,000; F1804, Sigma-Aldrich) and rabbit anti-β-tubulin antibody (1:6,000; ab6046, Abcam). For immunoblot detection, IRDye 800CW donkey anti-mouse IgG (926-32212; LI-COR Biosciences; 800-nm channel, displayed in green) and IRDye 680RD donkey anti-rabbit IgG (926-68073; LI-COR Biosciences; 700-nm channel, displayed in red) secondary antibodies were used. Immunoblot quantification in Suppl. Fig. 6 was performed by calculating the area under the curve using Fiji’s ‘Gel’ plugin.

### Live-cell imaging and analysis

For all live-cell imaging experiments, cells were seeded in eight-well chamber slides (µ-Slide, ibidi), and drugs were added directly to the imaging medium before acquisition. Where indicated, cells were stained with 50 nM SiR-DNA (Spirochrome), according to the manufacturer’s instructions, and/or treated with 150 nM paclitaxel (Sigma-Aldrich), 50 nM SiR-tubulin (Spirochrome), 10 µM DMA, or 0.33 µM nocodazole. The conditions used specifically for the microtubule stability assay are described below.

Mitotic timing was assessed by differential interference contrast (DIC) imaging using a Nikon Eclipse microscope (Nikon) equipped with a 20×/0.75 NA objective and a Flash 4.0 CMOS camera (Hamamatsu). CyclinA2–mEmerald fluorescence was quantified using a 40×/1.30 NA oil-immersion objective on the same microscope. Single-plane images were acquired every 3 min for 12–18 h using Micro-Manager (µManager).

Mitotic errors were assessed by time-lapse imaging of SiR-DNA-stained cells, primarily using the Nikon Eclipse microscope with a 40×/1.30 NA oil-immersion objective. Images were acquired every 3 min. Some experiments were performed using a Marianas spinning-disk confocal microscope system (Intelligent Imaging Innovations, Inc.) equipped with a 40× oil-immersion objective, 445-, 488-, 514-, and 561-nm lasers, a Flash 4.0 CMOS camera (Hamamatsu), and SlideBook 6 software (Intelligent Imaging Innovations, Inc.). Multinucleated cells were also imaged using the spinning-disk system. Micronuclei and DNA-damage markers were imaged using an Opera Phenix high-content imaging system (PerkinElmer).

For fixed-cell fluorescence analyses, nuclear Ki67 signal was quantified by applying DAPI-derived masks to the Ki67 channel, followed by background correction. Kinetochore-associated PLK1 was quantified using thresholded CENP-B masks applied to the PLK1 channel. Background fluorescence was measured in chromatin regions away from kinetochores and subtracted from the kinetochore-associated signal. Spindle length was measured in Fiji (ImageJ) as the pole-to-pole distance, and prometaphase cells were identified by the presence of MAD1-positive kinetochores.

### Quantification of CyclinA2–mEmerald fluorescence and degradation kinetics

For each cell, the mean CyclinA2–mEmerald fluorescence intensity was measured within a region of interest encompassing the cell using Fiji (ImageJ). The mean background intensity was measured in a cell-free region of the same image and subtracted from the mean cellular intensity. Nuclear envelope breakdown (NEBD) was identified by DIC imaging and defined as time zero (t = 0). Background-corrected fluorescence values were normalized to the value at NEBD (set to 100%) to generate individual-cell time courses. Local degradation rates were calculated using a five-point sliding window, and the maximum degradation rate was defined as the most negative local rate. The percentage of CyclinA2–mEmerald fluorescence remaining at 21 min was calculated relative to the value at NEBD. A Matlab scrip was used to obtain the results and the graphs

### Immunofluorescence and image acquisition

For visualization with most antibodies, coverslips with cells were washed twice with 1X PBS and fixed and permeabilised for 20 min at room temperature in 4% formaldehyde and 0.1% Triton X-100 in 1× PBS. Primary antibodies were diluted in stock solutions of PBS-BSA containing 0.1% Tween-20. After fixation, samples were blocked for 30 min in 3% BSA and 0.1% Tween-20 in PBS. Coverslips were incubated with primary Ab for 1–3 hr at room temperature, washed for 2 × 10 min in and then incubated with secondary antibody for 45 minutes, washed twice for 10minutes with 0.1% Triton-PBS-BSA solution and stained DAPI (Invitrogen, H3570), (1 µg/ml) diluted in 0.1%Triton-PBS-BSA); when required, Phalloidin-FITC (Sigma, P5282), (1:10.000) was included in the same staining solution. Two subsequent 10 min washes were done followed by a 10 min wash in 1X PBS. Coverslips were mounted on slides using ProLong Gold antifade reagent (Molecular Probes). Following fixation, were washed twice with PBS and maintained in PBS until image acquisition using the Opera Phenix system. Multinucleated cells were also imaged using the spinning-disk system. Micronuclei and DNA-damage markers were imaged using an Opera Phenix high-content imaging system (PerkinElmer).

For fixed-cell fluorescence analyses, nuclear Ki67 signal was quantified by applying DAPI-derived masks to the Ki67 channel, followed by background correction. Kinetochore-associated PLK1 was quantified using thresholded CENP-B masks applied to the PLK1 channel. Background fluorescence was measured in chromatin regions away from kinetochores and subtracted from the kinetochore-associated signal. Spindle length was measured in Fiji (ImageJ) as the pole-to-pole distance, and prometaphase cells were identified by the presence of MAD1-positive kinetochores.

| **Antibody / target** | **Host** | **Supplier** | **Catalogue no.** | **Application** | **Dilution** |
|---|---|---|---|---|---|
| CyclinA2 (N-terminal) | Mouse | Moravian Biotechnology | Custom-made | WB | 1:2000 |
| β-Tubulin | Rabbit | Abcam | ab6046 | WB and IF | 1:6000 WB and 1:2000 IF |
| Pericentrin | Rabbit | Abcam | ab4448 | IF | 1:1000 |
| MAD1 (clone 9B10) | Mouse | Sigma-Aldrich, 2 mg/ml | M8069 | IF | 1:200 |
| γH2AX | Rabbit | Sigma | 07627 | IF | 1:1000 |
| PLK1 phospho-Thr210 | Mouse | Abcam | Clone 2-A3 ab39068 | IF | 1:500 |
| Ki67 | Rabbit | Abcam | ab15580 | IF | 1:2000 |
| FLAG M2 | Mouse monoclonal | Sigma-Aldrich | F1804 | WB | 1:6000 |

### Mitotic-cell enrichment for mass spectrometry

Parental CyclinA2-mEmerald RPE1 cells (clone 6E) and the ΔD2ΔABBA mutant clones 3F and 10C were seeded in 15-cm dishes at 1 × 10⁶ cells per dish in 18 ml of medium. For each cell line, 30 dishes were prepared and divided into three biological replicates of 10 dishes each; 29 dishes were used for clone 10C, with nine dishes in the first replicate. Cells were treated with 150 nM taxol approximately 4 h 35 min before collection. Mitotic cells were harvested by adding approximately 1 ml of ice-cold PBS to each dish and gently shaking to detach the rounded mitotic cells while leaving the remaining cells attached. The procedure was repeated until the majority of mitotic cells had detached, and cells from the dishes assigned to each replicate were pooled. A separate aliquot, corresponding to one tenth of each sample, was collected for Western blotting. Remaining non-mitotic cells were collected from one dish per replicate for use as controls. Cell suspensions were washed twice, pelleted at 1300 rpm for 5 min, frozen on dry ice and stored at −80°C.

### TMT-based quantitative proteomics and phosphoproteomics

Cell pellets were lysed in a buffer containing 1% sodium deoxycholate (SDC), 100 mM triethylammonium bicarbonate (TEAB), 10% isopropanol and 50 mM NaCl, freshly supplemented with 5 mM TCEP (Thermo, Bond-breaker), 10 mM iodoacetamide (IAA), universal nuclease 1:2000 vol/vol (Pierce, #88700) and Halt protease and phosphatase inhibitor cocktail (Thermo, #78442, 100X) with 5 min of bath sonication. Protein concentration was measured with the Quick Start Bradford protein assay (Bio-Rad). Aliquots of 300 μg of total protein were digested overnight with trypsin (Pierce, 1:40) at room temperature. Peptides were labelled using nine labels of the TMT-10plex reagents (Thermo) by adding 40 μL of the reagent (20 μg/μL) into 100 μL of sample volume. The TMT-labelled peptide mixture was acidified with formic acid at 2% and the precipitated SDC was removed by centrifugation. The peptide pool was fractionated with high pH Reversed-Phase chromatography using the XBridge C18 column (2.1 × 150 mm, 3.5 μm, Waters) on an UltiMate 3000 HPLC system over a 1% gradient in 35 min. Mobile phase A was 0.1% (v/v) ammonium hydroxide and mobile phase B was 0.1% ammonium hydroxide (v/v) in acetonitrile.

Phosphopeptide enrichment of 24 peptide fractions was performed with the High-Select™ Fe-NTA Phosphopeptide Enrichment resin (Thermo) using a modified protocol in a well plate array format. A volume of 50 μL of resin/buffer was transferred on top of 10 μL filter tips that were fitted on a 96-well plate using a suitable tip rack. The resin was washed three times with 40 μL wash/binding solution and centrifugation at 500 g for 1 min. Peptides were reconstituted in 30 μL wash/binding solution and were loaded onto the tip-columns with the resin. After 30 min, the flow-through from three washes with wash/binding solution were collected in a clean 96-well plate with centrifugation at 500 g for 1 min each time. Phosphopeptides were eluted twice with 40 μL elution buffer in a clean 96-well plate with centrifugation at 500 g for 1 min and SpeedVac dried.

LC-MS analysis was performed on a Vanquish Neo UHPLC system coupled to the Orbitrap Ascend mass spectrometer (Thermo) using a 25 cm capillary column (Waters, nanoE MZ PST BEH130 C18, 1.7 μm, 75 μm × 250 mm) over a 110 min (proteome) or 80 min (phosphoproteome) gradient 5%-35% of mobile phase B composed of 80% acetonitrile, 0.1% formic acid. Peptides were preconcentrated onto a PEPMAP 100 C18 5μm 0.3 × 5 mm 1500 bar trapping column, and the analytical column was connected to a stainless-steel emitter on the Nanospray Flex ion source. For the proteome analysis (IMAC flowthrough), MS spectra were collected at Orbitrap mass resolution of 120,000 and precursors were selected for HCD fragmentation in the top speed mode (3 sec) with collision energy 32% and iontrap detection in turbo scan rate. MS3 scans were triggered by Real Time Search (RTS) against a fasta file containing UniProt Homo sapiens reviewed canonical and isoform sequences with multi-notch isolation (10 notches) and HCD fragmentation with collision energy 65% at 45,000 Orbitrap resolution. Selected precursors were dynamically excluded from further activation for 45 seconds with 10 ppm mass tolerance and RTS close-out was enabled with max 4 peptides per protein. Static modifications for RTS were TMT6plex at K/n-term (+229.1629), Carbamidomethyl at C (+57.0215) and variable modifications were Deamidated NQ (+0.984) and Oxidation of M (+15.9949) with maximum 1 missed-cleavage and 2 variable modifications per peptide. The phosphopeptide enriched fractions were analysed with an HCD MS2 method with collision energy 36% at 45,000 Orbitrap resolution.

The Sequest HT and Comet nodes in Proteome Discoverer 3.0 (Thermo) were used to search the raw mass spectra against a fasta file containing reviewed UniProt Homo sapiens entries. The precursor mass tolerance was set at 20 ppm and the fragment ion mass tolerance at 0.5 Da (or 1 Da for Comet) for the proteome data, and 0.02 Da for the phosphoproteome data, with up to 2 trypsin missed-cleavages allowed. TMT6plex at N-terminus/K and Carbamidomethyl at C were defined as static modifications. Dynamic modifications were oxidation of M and deamidation of N/Q, as well as phosphorylation of S/T/Y for the phosphoproteome data. Peptide confidence was estimated with the Percolator node and peptide FDR was set at 0.01 based on target-decoy search. Only unique peptides were used for quantification, considering protein groups for peptide uniqueness. Peptides with average reporter signal-to-noise greater than 3 were used for protein quantification. Phosphorylation localization was estimated with the IMP-ptmRS node.

### Spindle length and mitotic-stage analysis

Cells were seeded onto round coverslips placed in 6-well plates, processed as described in the “Immunofluorescence and image processing” section. They were stained with rabbit anti-pericentrin and mouse anti-MAD1 antibodies and imaged using a Marianas confocal spinning-disk microscope with a 60× oil-immersion objective. MAD1 staining was used to distinguish prometaphase from metaphase cells. Spindle length was calculated as the three-dimensional distance between the two Pericentrin-labelled centrosomes using the 3D ImageJ Suite plugin in ImageJ ^49^.

### Microtubule stability assay

The assay was adapted from a published kinetochore-microtubule depolymerization assay^50^. Cells were labelled with 20 nM SiR-tubulin, and microtubule depolymerization was induced with 0.66 µM nocodazole. Images were acquired every 30 s for 20 min using a spinning-disk confocal microscope equipped with a 60× oil-immersion objective. Individual-cell SiR-tubulin fluorescence traces were extracted using Fiji, normalized to the first time point (t = 0, set to 100%), and pooled from three independent experiments. In a complementary analysis, exact duplicate profiles were excluded, and the observed normalized traces were compared over the common 30-s to 15-min interval using a two-way repeated-measures ANOVA with Greenhouse–Geisser correction. AUC from 0 to 900 s was calculated by trapezoidal integration and compared between genotypes using Welch’s one-way ANOVA, followed by Holm-adjusted pairwise Welch’s t-tests. Curve fitting was performed in MATLAB; statistical analyses were performed in Python 3.12.14 using SciPy 1.17.0 and custom scripts.

### Cell competition assay

For cell competition assays, 25,000 untagged parental RPE1 cells were mixed with 25,000 emiRFP670nano3-positive CyclinA2–mEmerald parental 6E cells, ΔABBA single-mutant clones (2C, 7G and 3H) or ΔD2ΔABBA double-mutant clones (10C and 3F) and seeded in 6-well plates. A sample was analysed by flow cytometry on day 0 to verify the initial proportion of the two cell populations. Every 2 days, cells were harvested for flow-cytometric analysis using a BD FACSymphony A5 Cell Analyzer and passaged by reseeding 50,000 cells per well. The proportion of emiRFP670nano3-positive cells was monitored over a 12-day period.

### Chromosome spreads

Cells were treated for 3 h with 100 ng ml^−1^ colcemid (GIBCO), trypsinized and recovered in a Falcon tube. The cell suspension was centrifuged for 3 min at 250 × g and resuspended in 5 ml of 75 mM KCl, added dropwise. After a 15 min incubation at 37 °C, ten drops of Carnoys Fixative (3:1 methanol:acetic acid) were added. Following a 5 min centrifugation at 200 × g, the cell pellet was resuspended in 5 ml of Carnoys Fixative. After 90 min at −20 °C, a second fixation was performed using 5 ml of Carnoys Fixative at room temperature for 15 min. Cells were centrifuged at 200 × g for 5 min, the supernatant removed, and the pellet resuspended in 200 µl of Carnoys Fixative. Spreads were performed by dropping the cells from a height of 30 to 40 cm onto wet slides in a wet chamber. Spreads were mounted and stained using Vectashied with DAPI (ReactoLab). Samples were imaged on a GE wide-field DeltaVision Elite microscope equipped with a 63x/1.2 NA Oil Objective and SoftWoRx Imaging software. The number of chromosomes per cell was counted using ImageJ software.

### DNA fibre assay

Exponentially growing RPE1 cells were sequentially labelled with iododeoxyuridine (IdU; [25 µM]) and chlorodeoxyuridine (CldU; 125 µM) for 20 min each. Cells were washed with ice-cold PBS and resuspended at 7.5 × 10⁵ cells/ml in cold PBS. A 2-µl aliquot of the cell suspension was deposited onto a glass slide and allowed to partially air-dry for 5 min. Cells were lysed by adding 7 µl of lysis buffer (200 mM Tris–HCl, pH 7.5, 50 mM EDTA and 0.5% SDS) and incubating for 2 min. Slides were tilted at 15° to spread the DNA fibres and air-dried horizontally. DNA fibres were fixed in methanol/acetic acid (3:1) for 10 min, washed in distilled water and denatured in 2.5 M HCl for 80 min. Following three 5-min washes in PBS, slides were blocked in 5% BSA in PBS for 25 min. Incorporated IdU and CldU were detected using mouse anti-BrdU (1:20) and rat anti-BrdU (1:400) antibodies, respectively, diluted in blocking buffer and incubated for 2 h in a humidified chamber. Slides were washed three times in PBS for 5 min and incubated with Cy3-conjugated sheep anti-mouse (1:500) and Alexa Fluor 488-conjugated goat anti-rat (1:400) secondary antibodies for 1 h, protected from light. Following three additional 5-min washes in PBS, slides were mounted using Vectashield mounting medium. Images were acquired using a Leica SP8 confocal microscope with a 63×/1.40 NA oil-immersion objective. Images were processed using Fiji (ImageJ 2.14.0/1.54f).

### Statistical analysis

For the siR-Tubulin decay assay, normalized single-cell fluorescence traces from three independent experiments were analysed over the common 0–900-s interval. Differences in fluorescence decay over time were assessed using a two-way repeated-measures ANOVA, with time as the within-cell factor and genotype as the between-cell factor, applying the Greenhouse–Geisser correction. Where indicated, each mutant was compared with the parental cells at individual time points using two-sided Welch’s t-tests with Šidák correction for the two comparisons performed at each time point. AUC₀–₉₀₀ s was calculated for each cell by trapezoidal integration. AUC distributions were compared using a Kruskal–Wallis test, followed by pairwise two-sided Mann–Whitney U tests with Holm correction across all three pairwise comparisons. Time-course data are presented as mean ± SEM; AUC plots show individual cells with the median. Adjusted P values are reported, with P < 0.05 considered statistically significant. Statistical analyses were performed in Python 3.12.14 using SciPy 1.17.0 and custom scripts. Local five-point degradation rates, time to maximum degradation rate, and normalised fluorescence remaining 21 min after NEBD were analysed separately for each treatment using ordinary one-way ANOVA followed by Dunnett’s multiple-comparisons test against the matched parental 6E control. Mitotic timing and IF measurements were analysed by one-way ANOVA followed by Dunnett’s multiple-comparisons test, comparing each mutant with the parental control. Ki67 IF measurements were averaged per biological experiment before statistical analysis. Statistical analyses of the CyclinA2 degradation curves and IF measurements were performed using GraphPad Prism.

## Supporting information

Supplemental FiguresS1-5

## Acknowledgements

We are grateful to all the members of the Cell Division Team, particularly Dr Luca Cirillo, for their comments and support during the genesis of this work. We also thank the Short Linear Motifs team for their insightful discussions, particularly Dr. Norman Davey for his invaluable support. We gratefully acknowledge the support provided by the ICR core facilities, particularly the Light Microscopy and Flow Cytometry facilities. This work was supported by Cancer Research UK (CRUK) Programme Grant C29/A24397 awarded to JP. SGS was supported by an Erasmus fellowship, and YW was supported by a summer studentship from the ICR.

## Author Contributions

**Annalisa Roberti**: Conceptualisation; Formal analysis; Supervision; Validation; Investigation; Methodology; Writing, original draft; Writing – review and editing. **Sofia Garcia Soto**: Investigation. **Ynon Weiss**: Investigation. **Theodoros I. Roumeliotis**: Formal analysis; Investigation; Methodology. **Jyoti S. Choudhary**: Formal analysis; Investigation; Methodology. **Jonathon Pines**: Conceptualisation; Formal analysis; Supervision; Funding acquisition; Validation; Writing—original draft; Project administration; Writing—review and editing.

The authors declare that they have no conflict of interest.

## References

1. Musacchio, A. & Salmon, E. D. The spindle-assembly checkpoint in space and time. Nat. Rev. Mol. Cell Biol. 8, 379–393 (2007).

2. Musacchio, A. Spindle assembly checkpoint: the third decade. Philos. Trans. R. Soc. Lond., B, Biol. Sci. 366, 3595–3604 (2011).

3. McAinsh, A. D. & Kops, G. J. P. L. Principles and dynamics of spindle assembly checkpoint signalling. Nat. Rev. Mol. Cell Biol. 24, 543–559 (2023).

4. Lara-Gonzalez, P., Pines, J. & Desai, A. Spindle assembly checkpoint activation and silencing at kinetochores. Semin. Cell Dev. Biol. 117, 86–98 (2021).

5. Barford, D. Understanding the structural basis for controlling chromosome division. Philos Trans A Math Phys Eng Sci 373, 20130392–20130392 (2015).

6. Yamano, H. APC/C: current understanding and future perspectives. F1000Research 8, F1000 Faculty Rev-725 (2019).

7. Pines, J. Cubism and the cell cycle: the many faces of the APC/C. Nat. Rev. Mol. Cell Biol. 12, 427–438 (2011).

8. Sudakin, V., Chan, G. K. T. & Yen, T. J. Checkpoint inhibition of the APC/C in HeLa cells is mediated by a complex of BUBR1, BUB3, CDC20, and MAD2. J. Cell Biol. 154, 925–936 (2001).

9. Alfieri, C., Chang, L., Zhang, Z., Yang, J., Maslen, S., Skehel, M. & Barford, D. Molecular basis of APC/C regulation by the spindle assembly checkpoint. Nature 536, 431–436 (2016).

10. Izawa, D. & Pines, J. The mitotic checkpoint complex binds a second CDC20 to inhibit active APC/C. Nature 517, 631–634 (2014).

11. Pines, J. & Hunter, T. Human cyclin A is adenovirus E1A-associated protein p60 and behaves differently from cyclin B. Nature 346, 760–3 (1990).

12. Coverley, D., Laman, H. & Laskey, R. A. Distinct roles for cyclins E and A during DNA replication complex assembly and activation. Nat. Cell Biol. 4, 523–528 (2002).

13. Furuno, N., Elzen, N. den & Pines, J. Human Cyclin A Is Required for Mitosis until Mid Prophase. J. Cell Biol. 147, 295–306 (1999).

14. Hégarat, N., Crncec, A., Rodriguez, M. F. S. P., Iturra, F. E., Gu, Y., Busby, O., Lang, P. F., Barr, A. R., Bakal, C., Kanemaki, M. T., Lamond, A. I., Novak, B., Ly, T. & Hochegger, H. Cyclin A triggers Mitosis either via the Greatwall kinase pathway or Cyclin B. EMBO J. 39, EMBJ2020104419 (2020).

15. Gong, D., Pomerening, J. R., Myers, J. W., Gustavsson, C., Jones, J. T., Hahn, A. T., Meyer, T. & Ferrell, J. E. J. Cyclin A2 regulates nuclear-envelope breakdown and the nuclear accumulation of cyclin B1. Curr. Biol. 17, 85–91 (2007).

16. Gong, D. & Ferrell, J. E. J. The Roles of Cyclin A2, B1, and B2 in Early and Late Mitotic Events. Mol. Biol. Cell (2010). doi:10.1091/mbc.e10-05-0393

17. Elzen, N. den & Pines, J. Cyclin A is destroyed in prometaphase and can delay chromosome alignment and anaphase. J. Cell Biol. 153, 121–136 (2001).

18. Geley, S., Kramer, E., Gieffers, C., Gannon, J., Peters, J. M. & Hunt, T. APC/C-dependent proteolysis of human cyclin A starts at the beginning of mitosis and is not subject to the spindle assembly checkpoint. J. Cell Biol. 153, 137–48. (2001).

19. Kaspar, M., Dienemann, A., Schulze, C. & Sprenger, F. Mitotic degradation of cyclin A is mediated by multiple and novel destruction signals. Curr Biol 11, 685–690 (2001).

20. Fiore, B. D. & Pines, J. How cyclin A destruction escapes the spindle assembly checkpoint. J. Cell Biol. 190, 501–509 (2010).

21. Fiore, B. D., Davey, N. E., Hagting, A., Izawa, D., Mansfeld, J., Gibson, T. J. & Pines, J. The ABBA Motif Binds APC/C Activators and Is Shared by APC/C Substrates and Regulators. Dev. Cell 32, 358–372 (2015).

22. Wolthuis, R., Clay-Farrace, L., Zon, W. van, Yekezare, M., Koop, L., Ogink, J., Medema, R. & Pines, J. Cdc20 and Cks direct the spindle checkpoint-independent destruction of cyclin A. Mol. Cell 30, 290–302 (2008).

23. Zhang, S., Tischer, T. & Barford, D. Cyclin A2 degradation during the spindle assembly checkpoint requires multiple binding modes to the APC/C. Nat. Commun. 10, 3863 (2019).

24. Evans, T., Rosenthal, E. T., Youngblom, J., Distel, D. & Hunt, T. Cyclin: A protein specified by maternal mRNA in sea urchin eggs that is destroyed at each cleavage division. Cell 33, 389–396 (1983).

25. Swenson, K., Farrell, K. M. & Ruderman, J. V. The clam embryo protein cyclin A induces entry into M-phase and the resumption of meiosis in Xenopus oocytes. Cell 47, 861–870 (1986).

26. Lehner, C. F. & O’Farrell, P. H. Expression and function of Drosophila cyclin A during embryonic cell cycle progression. Cell 56, 957–968 (1989).

27. Hunt, T., Luca, F. C. & Ruderman, J. V. The requirements for protein synthesis and degradation, and the control of destruction of cyclins A and B in the meiotic and mitotic cell cycles of the clam embryo. J. Cell Biol. 116, 707–724 (1992).

28. Luca, F. C. & Ruderman, J. V. Control of programmed cyclin destruction in a cell-free system. J. Cell Biol. 109, 1895–1909 (1989).

29. Whitfield, W. G. F., Gonzalez, C., Maldonado, C. G. & Glover, D. M. The A- and B-type cyclins of Drosophila are accumulated and destroyed in temporally distinct events that define separable phases of the G2-M transition. EMBO J. 9, 2563–2572 (1990).

30. Minshull, J., Golsteyn, R., Hill, C. S. & Hunt, T. The A- and B-type cyclin associated cdc2 kinases in Xenopus turn on and off at different times in the cell cycle. EMBO J. 9, 2865–2875 (1990).

31. Kabeche, L. & Compton, D. A. Cyclin A regulates kinetochore microtubules to promote faithful chromosome segregation. Nature 502, 110–113 (2013).

32. Valles, S. Y., Bural, S., Godek, K. M. & Compton, D. A. Cyclin A/Cdk1 promotes chromosome alignment and timely mitotic progression. Mol. Biol. Cell 35, ar141 (2024).

33. Dumitru, A. M. G., Rusin, S. F., Clark, A. E. M., Kettenbach, A. N. & Compton, D. A. Cyclin A/Cdk1 modulates Plk1 activity in prometaphase to regulate kinetochore-microtubule attachment stability. eLife 6, e29303 (2017).

34. Crncec, A., Lau, H. W., Ng, L. Y., Ma, H. T., Mak, J. P. Y., Choi, H. F., Yeung, T. K. & Poon, R. Y. C. Plasticity of mitotic cyclins in promoting the G2–M transition. J. Cell Biol. 224, e202409219 (2025).

35. Örd, M., Winters, M. J., Subbanna, M. S., Garrido, N. de M., Cushing, V. I., Kliche, J., Benz, C., Ivarsson, Y., Greber, B. J., Pryciak, P. M. & Davey, N. E. High-throughput investigation of cyclin docking interactions reveals the complexity of motif binding determinants. Nat. Commun. 16, 7622 (2025).

36. Jinek, M., Chylinski, K., Fonfara, I., Hauer, M., Doudna, J. A. & Charpentier, E. A Programmable Dual-RNA–Guided DNA Endonuclease in Adaptive Bacterial Immunity. Science 337, 816–821 (2012).

37. Matlashov, M. E., Shcherbakova, D. M., Alvelid, J., Baloban, M., Pennacchietti, F., Shemetov, A. A., Testa, I. & Verkhusha, V. V. A set of monomeric near-infrared fluorescent proteins for multicolor imaging across scales. Nat. Commun. 11, 239 (2020).

38. Zerjatke, T., Gak, I. A., Kirova, D., Fuhrmann, M., Daniel, K., Gonciarz, M., Müller, D., Glauche, I. & Mansfeld, J. Quantitative Cell Cycle Analysis Based on an Endogenous All-in-One Reporter for Cell Tracking and Classification. Cell Rep. 19, 1953–1966 (2017).

39. Collin, P., Nashchekina, O., Walker, R. & Pines, J. The spindle assembly checkpoint works like a rheostat rather than a toggle switch. Nat. Cell Biol. 15, 1378–1385 (2013).

40. Glotzer, M., Murray, A. W. & Kirschner, M. W. Cyclin is degraded by the ubiquitin pathway. Nature 349, 132–138 (1991).

41. Pfleger, C. M. & Kirschner, M. W. The KEN box: an APC recognition signal distinct from the D box targeted by Cdh1. Genes & development 14, 655–665 (2000).

42. al-Rawi, A., Kaye, E., Korolchuk, S., Endicott, J. A. & Ly, T. Cyclin A and Cks1 promote kinase consensus switching to non-proline-directed CDK1 phosphorylation. Cell Rep. 42, 112139 (2023).

43. Cuylen, S., Blaukopf, C., Politi, A. Z., Müller-Reichert, T., Neumann, B., Poser, I., Ellenberg, J., Hyman, A. A. & Gerlich, D. W. Ki-67 acts as a biological surfactant to disperse mitotic chromosomes. Nature 535, 308–312 (2016).

44. Sanz-Flores, M., Ruiz-Torres, M., Aguirre-Portolés, C., Bakkali, A. E., Salvador-Barberó, B., Villarroya-Beltri, C., Ortega, S., Megías, D., Gerlich, D. W., Álvarez-Fernández, M. & Malumbres, M. PP2A-B55 phosphatase counteracts Ki-67-dependent chromosome individualization during mitosis. Cell Rep. 43, 114494 (2024).

45. Remnant, L., Kochanova, N. Y., Reid, C., Cisneros-Soberanis, F. & Earnshaw, W. C. The intrinsically disorderly story of Ki-67. Open Biol. 11, 210120 (2021).

46. Sobecki, M., Mrouj, K., Colinge, J., Gerbe, F., Jay, P., Krasinska, L., Dulic, V. & Fisher, D. Cell-Cycle Regulation Accounts for Variability in Ki-67 Expression Levels. Cancer Res. 77, 2722–2734 (2017).

47. Lanni, J. S. & Jacks, T. Characterization of the p53-Dependent Postmitotic Checkpoint following Spindle Disruption. Mol. Cell. Biol. 18, 1055–1064 (1998).

48. Stead, E., White, J., Faast, R., Conn, S., Goldstone, S., Rathjen, J., Dhingra, U., Rathjen, P., Walker, D. & Dalton, S. Pluripotent cell division cycles are driven by ectopic Cdk2, cyclin A/E and E2F activities. Oncogene 21, 8320–8333 (2002).

49. Ollion, J., Cochennec, J., Loll, F., Escudé, C. & Boudier, T. TANGO: a generic tool for high-throughput 3D image analysis for studying nuclear organization. Bioinformatics 29, 1840–1841 (2013).

50. Dudka, D., Castrogiovanni, C., Liaudet, N., Vassal, H. & Meraldi, P. Spindle-Length-Dependent HURP Localization Allows Centrosomes to Control Kinetochore-Fiber Plus-End Dynamics. Curr. Biol. 29, 3563–3578.e6 (2019).

