## Supplemental FiguresS1-5 for "Why is Cyclin A degraded before Cyclin B?"

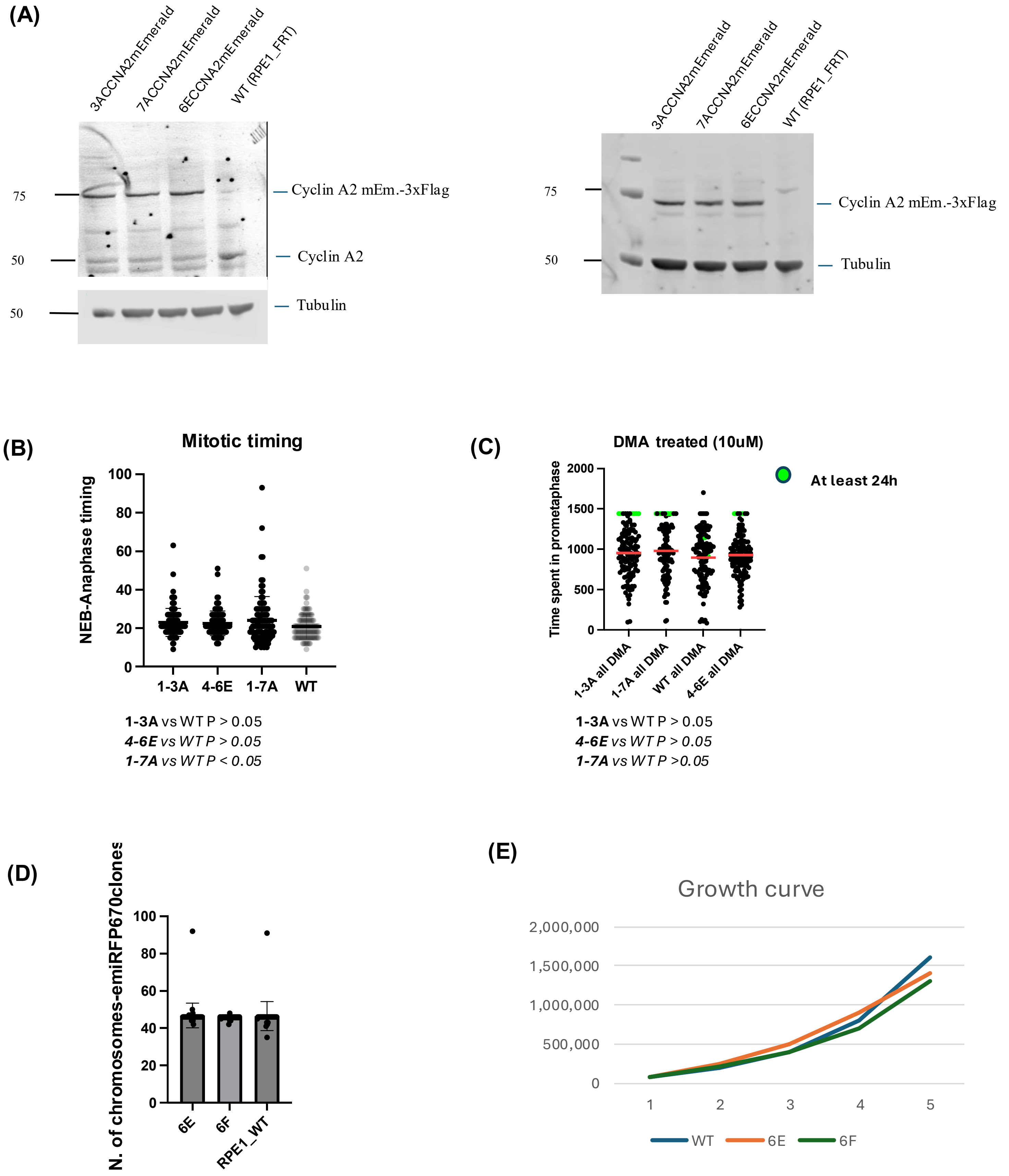


**Supplementary Figure S1. Cyclin A2 expression, mitotic timing, DMA treatment, chromosome number and cell growth.**

**(A)** Western blot analysis of Cyclin A2 and Flag-tagged Cyclin A2 in the indicated cell lines. Anti-Cyclin A blotted on the left, anti-Flag blotted on the right. Tubulin was used as a loading control. Molecular mass markers in kDa on the left. **(B)** Time from nuclear envelope breakdown (NEBD) to anaphase in clones 3A, 6E and 7A and WT cells. Each dot represents an individual cell. 100 cells analysed in 3 independent replicates. **(C)** Time spent in mitosis following treatment with 10 µM DMA. Each dot represents an individual cell. 100 cells analysed in 3 independent replicates. **(D)** Quantification of metaphase spreads in the indicated cell lines. >15 cells analysed in 2 independent replicates. **(E)** Growth curves used to compare cell doubling times. N=2.

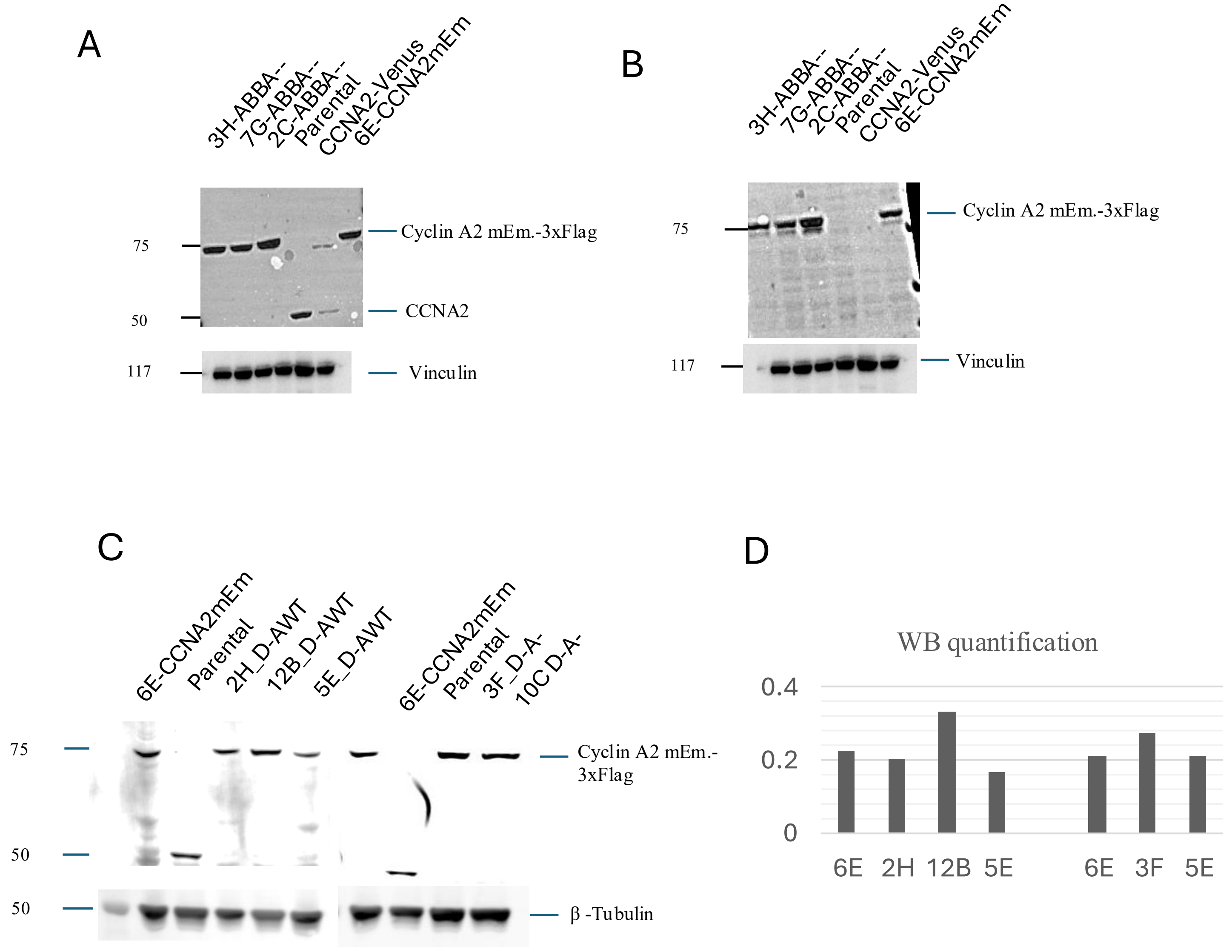


**Supplementary Figure S2**. Western blot analysis of Cyclin A2 in the indicated cell lines and clones. Representative immunoblots probed with antibodies against Cyclin A2 (panels A & C) and 3×Flag (panel B). Vinculin was used as a loading control in panels A and B and tubulin in panel C. Molecular mass markers in kDa on the left. Panels A - C show representative fluorescent western blots of the indicated parental and mutant clones. Panel D shows the quantification of CyclinA2 protein in the different ΔD2 and ΔD2box ΔABBA clones

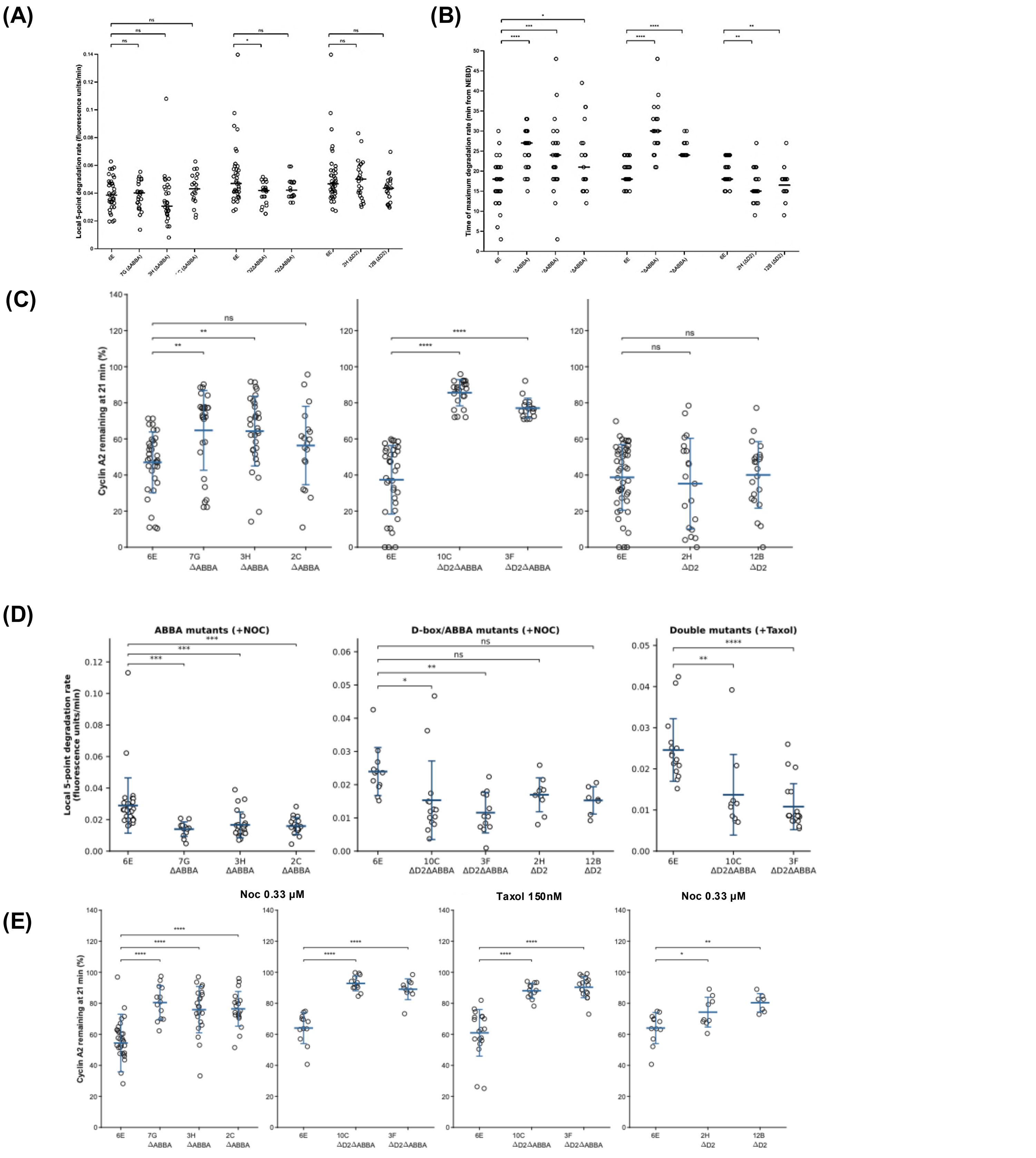


**Supplementary Figure S3. Quantification of Cyclin A2 degradation kinetics in unperturbed, nocodazole- and taxol-treated cells**

**(A)** Local max. five-point degradation rate **(B)** time to maximum degradation rate and **(C)** Cyclin A2 remaining at 21 min across the indicated cell lines in untreated conditions. Each dot represents an individual cell; data are shown as mean ± SD. At least 17 cells per clone were analysed, pooled from three independent experiments. **(D)** Local maximum five-point degradation rate in ABBA mutants, D2box/ABBA mutants (double mutants) and D2box mutants treated with 3.3 µm nocodazole (left and middle), or with 150nM taxol (right). **(E)** Cyclin A2 remaining 21 min after NEBD in ABBA mutants treated with (far left), double mutants treated with 3.3 µm nocodazole (left) or 50nM taxol (right), and D2-box mutants treated with 3.3 µm nocodazole (far right). Between 7 and 30 cells per clone were analysed from at least two independent experiments. Statistical comparisons are indicated above the plots and were performed against the corresponding parental 6E control using one-way ANOVA followed by Dunnett's multiple-comparisons test and are indicated above the plots. ns, $P \geq 0.05$; $^{*}P < 0.05$; $^{**}P < 0.01$; $^{***}P < 0.001$; $^{****}P < 0.0001$.

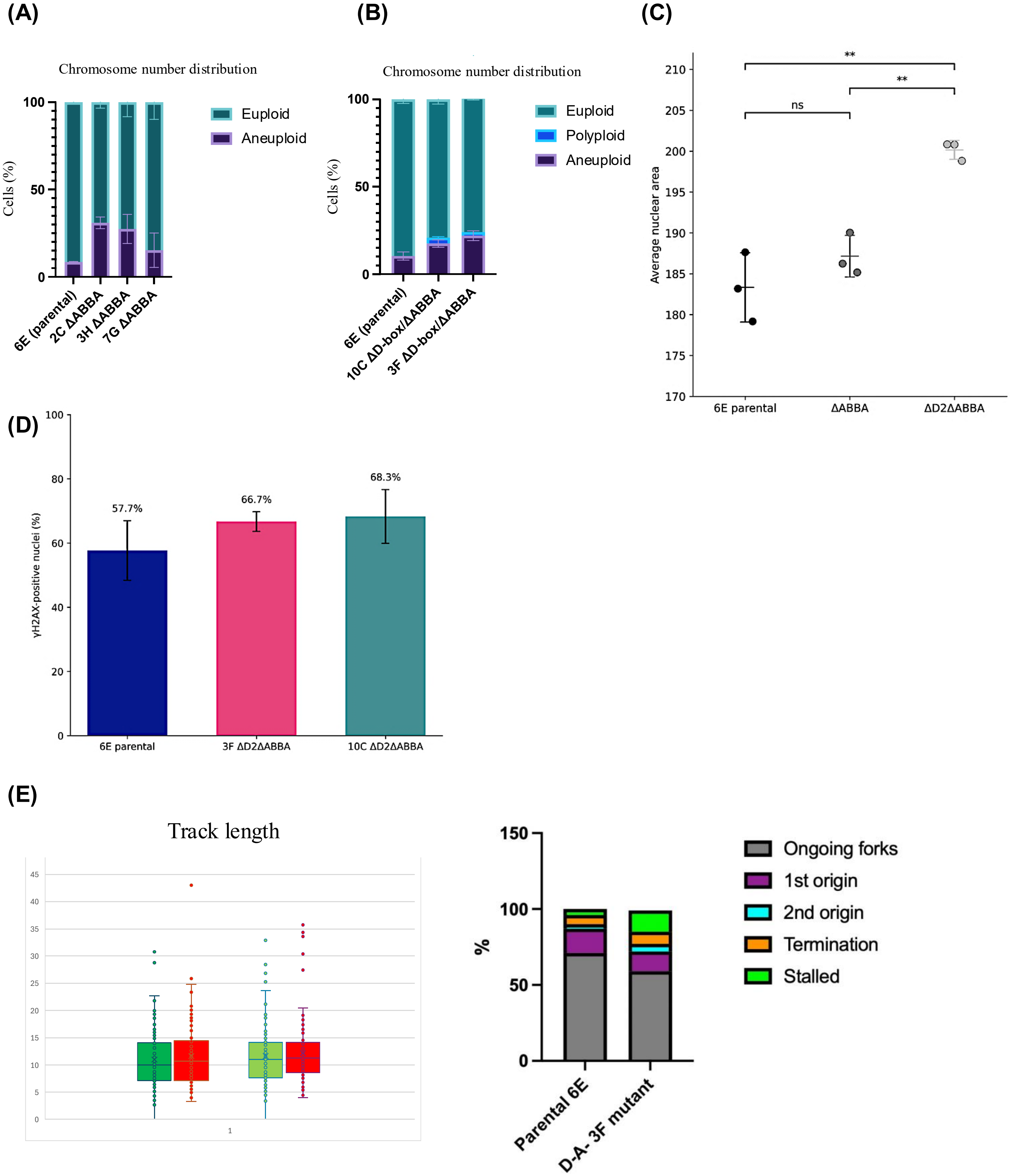


**Supplementary Figure S4. Chromosome number, nuclear area, DNA damage and replication-fork analysis in Cyclin A2 mutant cells.**
**(A)** Distribution of euploid and aneuploid metaphases in parental 6E and ΔABBA clones 2C, 3H and 7G. **(B)** Distribution of euploid, polyploid and aneuploid metaphases in parental 6E and D2box/ABBA mutant clones 10C and 3F. >15 cells analysed in 2 independent replicates. **(C)** Mean nuclear area in parental 6E, ΔABBA and ΔD2box ΔABBA cells; statistical comparisons are indicated above the plot. More than 2000 cells analysed from 2 independent experiments **(D)** Percentage of γH2AX-positive cells in parental 6E and ΔD2box ΔABBA mutant clones 3F and 10C. Values are labelled above the bars. At least 50 cells analysed/cell line from 3 independent experiments **(E)** DNA-fibre analysis comparing parental 6E and the ΔD2box ΔABBA mutant 3F: replication-track length (left) and the proportions of ongoing forks, first-origin firing, second-origin firing, termination and stalled forks (right).

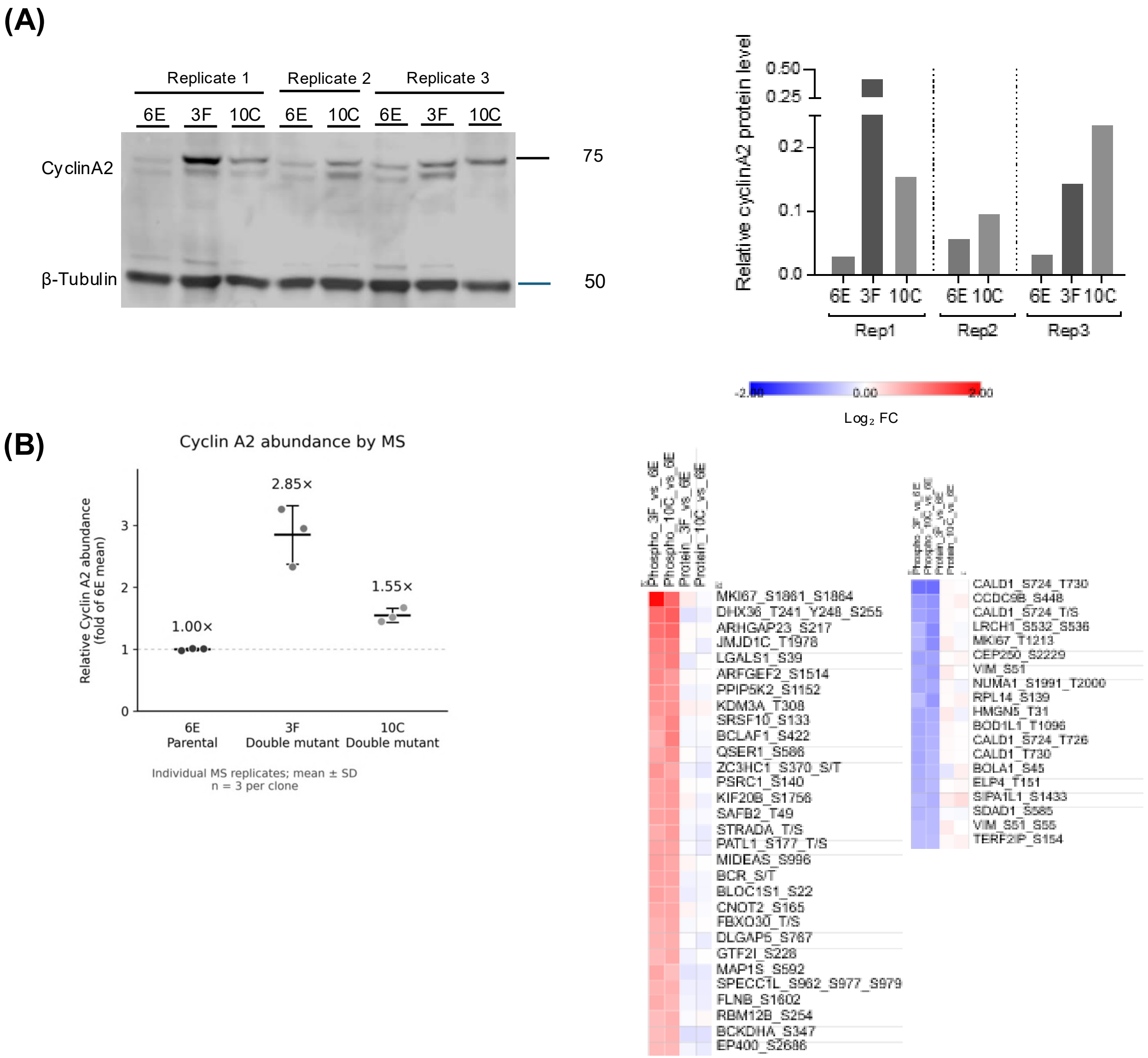


**Supplementary Figure S5. Mitotic enrichment and sample preparation for phospho-proteomic analysis.** **(A)** Representative immunoblot (left) and quantification of Cyclin A2 (right) in samples from three replicates of cultures enriched in mitosis with 150 nM taxol. The molecular mass markers are shown in kDa. These samples were submitted for phospho-proteomic mass spectrometry (MS). The 3F ΔABBA/ΔD2box mutant sample from replicate 2 is missing from the immunoblot but was included in the mass spectrometry analysis. **(B)** Relative Cyclin A2 abundance measured by TMT-based mass spectrometry in parental (6E) and ΔABBA/ΔD2box mutant (3F and 10C) clones. Normalized protein abundances were divided by the mean abundance in parental cells. Each dot represents an individual replicate (n = 3 per clone); horizontal lines and error bars indicate mean ± SD. Numbers above the groups indicate mean fold changes relative to the parental control.
**(C)** Heatmaps showing $\log_2$ fold changes in phospho-peptide abundance and corresponding total protein abundance in Cyclin A2 mutant clones 3F and 10C relative to the parental clone 6E, as determined by quantitative mass spectrometry. Phospho-peptides are grouped according to their concordant increase (left) or decrease (right) in both mutant clones. Each row represents an individual phospho-peptide, with the first two columns showing phospho-peptide abundance changes and the last two columns showing the corresponding total protein abundance changes. The corresponding quantitative data are provided in Supplementary Table S1.
